# Mass Transfer Formulations Shape Oxygen Transport and Tissue Oxygenation in Microvascular Networks

**DOI:** 10.64898/2026.11.19.767880

**Authors:** Gaia Stievano, Dominic Campbell-Pitt, Giorgia Barra, Timo Koch, Franca Schmid

## Abstract

**Purpose:** Continuous oxygen delivery is essential for cerebral tissue function, yet its local supply characteristics remain incompletely understood. Quantifying oxygen transport in microvascular networks (MVNs) requires a physiologically consistent description of oxygen exchange between vascular compartments via mass transfer coefficients (MTCs). We investigate how different MTC formulations influence oxygen availability and hypoxic burden in realistic MVNs, and whether they produce relevant local differences in tissue oxygenation

**Methods:** A multicompartment oxygen transport model describing exchange between red blood cells (RBCs), plasma, and tissue in realistic MVNs was developed and evaluated against in vivo measurements. Three RBC–plasma MTC formulations were compared: constant, hematocrit-dependent, and hematocrit- and saturation-dependent. Additionally, a double-flux formulation incorporating a direct RBC-to-tissue flux was compared with standard single-flux formulations.

**Results:** Global tissue oxygen distributions were comparable across RBC–plasma MTC formulations, but hematocrit- and saturation-dependent MTC formulations predicted larger hypoxic tissue fractions. The largest differences occurred under low hematocrit, with local plasma oxygen partial pressure 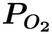 differing by ∼6 mmHg along representative vascular paths. Introducing the direct RBC–tissue exchange reduced tissue 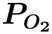 at low hematocrit and enhanced it at high hematocrit, with the largest differences observed in small vessels. Sensitivity analysis revealed that RBC oxygen diffusivity can alter the double-flux effect magnitude.

**Conclusion:** MTC assumptions substantially influence predicted oxygen availability and hypoxic burden, particularly under low-hematocrit conditions and in small vessels, underlining the need for their accurate definition. The proposed model provides a basis for evaluating these effects, improving the interpretation of oxygen transport predictions in realistic MVNs.

## 1 Introduction

The cerebral microcirculation plays a central role in delivering nutrients and removing metabolic waste products [1, 2]. Within these processes, oxygen transport is particularly critical for maintaining cerebral homeostasis, as oxygen is a fundamental driver of cellular energetics [3]. Consequently, to understand both healthy brain function and impairments during neurological disorders, it is essential to characterize transport processes at the microvascular scale.

In arterial blood, approximately 98% of oxygen is reversibly bound to hemoglobin within red blood cells (RBCs), while the remaining fraction is dissolved in plasma and in the hemoglobin solution inside RBCs [4]. During oxygen extraction, oxygen dissociates from hemoglobin and diffuses across the RBC membrane, plasma, and capillary wall before reaching parenchymal cells, where it is consumed by mitochondria. Maintaining adequate oxygenation therefore relies on efficient transport from the vascular system to surrounding tissue. This process is particularly critical in the brain, where energy storage is limited and neural cells depend on continuous oxygen delivery [5]. This raises the question of whether microvascular impairments, as observed during aging and neurodegenerative diseases, do indeed have the potential to induce localized hypoxia and thereby contribute to disease progression [6–8]. As such, oxygen transport could provide an underlying mechanistic explanation for why vascular integrity is increasingly recognized as a key determinant of brain health [9–13].

Quantitative characterization of oxygen transport in the cerebral microcirculation remains challenging due to its complex vascular architecture, which is characterized by small spatial scales, heterogeneous perfusion, and dynamically varying metabolic demands [14–19]. While microscale measurements of oxygenation remain prohibitive in humans, two-photon phosphorescence lifetime imaging in mice provides a useful technique to assess vascular and tissue oxygenation with high spatial resolution. However, the experimental complexity limits sample size and requires advanced imaging setups if deeper brain tissues are to be investigated [20–22]. This motivates the use of in silico models, which enable controlled investigation of oxygen transport under different conditions.

Computational modeling of oxygen transport from the microvasculature to the surrounding tissue was pioneered by the Krogh model (1919) [23]. However, its simplifying assumptions regarding vascular geometry and hemodynamics limit its ability to capture the complexity of real microvascular systems. More advanced approaches have since been developed to simulate oxygen transport in reconstructed or synthetic vascular networks consisting of up to 256,000 vessels, while accounting for blood flow dynamics and heterogeneous oxygen extraction [24–33]. Additionally, models resolving oxygen transport at the scale of individual RBCs have been developed to investigate the mechanisms governing blood–tissue oxygen exchange [34–37]. While the latter are restricted to very small simulation domains, the former network-scale models rely on several assumptions that may affect the accuracy of predicted oxygen distributions.

A common assumption in current oxygen transport models is the use of a single blood-phase oxygen partial pressure (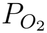), implying that blood plasma and RBCs have identical oxygen levels. This simplification neglects oxygen gradients that arise between RBCs and plasma, commonly referred to as erythrocyte-associated transients (EATs) [18, 38–43]. As a result, many models disregard the resistance to oxygen exchange between RBCs and plasma and consider oxygen exchange only between plasma and tissue [24, 31, 32, 44–49]. Experimental evidence, however, indicates that both 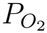 and EATs vary along the vessel, depending on local hematocrit, RBC spacing, flow conditions, and tissue oxygen demand [18, 34–37, 40, 50–52].

To address this limitation, we develop a mixed-dimensional oxygen transport model in which microvessels are represented as one-dimensional edges embedded within a three-dimensional tissue domain. The model describes oxygen transport in three compartments: RBCs, plasma, and tissue and is applicable to realistic microvascular networks (MVNs). The underlying blood flow model accounts for vessel-specific hematocrit variations. To describe oxygen exchange between the vasculature and tissue, two strategies are considered. In the *Single-Flux Model*, oxygen is transported sequentially from RBCs to plasma and from plasma to tissue, driven by the respective 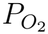 gradients and limited by the corresponding transport resistances. In the *Double-Flux Model*, an additional direct pathway from RBCs to tissue is introduced. This pathway becomes relevant when capillaries and RBCs are comparable in size (i.e., *≤* 6 *µ*m), such that RBCs can come into direct contact with the vessel wall. Under these conditions, both theoretical studies and in vivo experiments have shown that tissue oxygenation fluctuates with the passage of individual RBCs [4, 38, 41, 53], suggesting that oxygen delivery may be influenced by the proximity of RBCs to the capillary wall and raising the question of whether a single oxygen flux is sufficient to describe capillary oxygen exchange.

In continuum-based oxygen transport models, including the one presented here, oxygen exchange between RBCs and plasma is described by a lumped mass transfer coefficient (*K*_cell_) that incorporates multiple transport resistances, including intracellular diffusion and oxygen transport across the RBC membrane, without explicitly resolving transport within individual RBCs. Previous studies suggest that *K*_cell_ depends on several physiological factors, including blood flow conditions, tissue oxygen demand, oxygen saturation, hematocrit, and RBC velocity [18, 34–37, 40, 50–52]. In particular, the dependence of *K*_cell_ on hematocrit is related to the spatial organization of RBCs within the vessel: increasing hematocrit reduces the plasma gaps between adjacent cells, shortening the diffusion path for oxygen and thereby enhancing oxygen transfer [54]. Its dependence on oxygen saturation reflects the contribution of facilitated diffusion through hemoglobin binding. At high saturation levels, the flattening of the hemoglobin–oxygen dissociation curve [55] reduces oxygen exchange associated with reversible binding, whereas intermediate saturation levels promote larger saturation changes for a given variation in 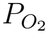, resulting in higher effective mass transfer coefficient (MTC) values. Despite this knowledge, the complex interplay of these mechanisms makes the experimental characterization of *K*_cell_ challenging. Consequently, many oxygen transport models adopt a constant value for *K*_cell_, likely limiting the accuracy of predicted oxygen distributions. In context of the *Single-Flux Model*, we assess the impact of this assumption by comparing a constant coefficient with literature-based formulations in which *K*_cell_ varies with hematocrit and oxygen saturation.

Taken together, we introduce a novel model for microvascular oxygen transport applicable to realistic MVNs that explicitly accounts for oxygen exchange between RBCs, plasma, and tissue. By systematically evaluating alternative descriptions of oxygen exchange and the associated MTCs, we quantify how physiological assumptions and parameter uncertainties influence predicted oxygen distributions. Thereby, we identify the mechanisms and parameters most strongly affecting microvascular oxygen delivery, highlighting the need for improved experimental characterization to accurately predict hypoxic tissue volumes and assess the robustness of cerebral oxygen supply.

## 2 Materials and Methods

### 2.1 Microvascular Hemodynamics

In the microcirculation, the Reynolds (Re *≪*1), Womersley (Wo *≪*1), and Dean (De*≪* 1) numbers are all very small, indicating laminar, quasi-steady flow with negligible curvature effects. These conditions justify a Poiseuille-type description of blood flow in the microcirculation [15, 19, 56–60]. Under the additional assumptions of negligible radial velocities, slender vessels (length much larger than radius), and rigid walls, the governing equations reduce to a one-dimensional description obtained by cross-sectional averaging. To this end, we describe each vessel segment by a parametrization of its centerline with the local variable *s ∈* [0, 1] and its cross-sectional area 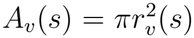, where *r*_*v*_ is the vessel radius (Figure 1(a)). A local cylindrical coordinate system (*r, θ, z*) is adopted.

**Fig. 1.**
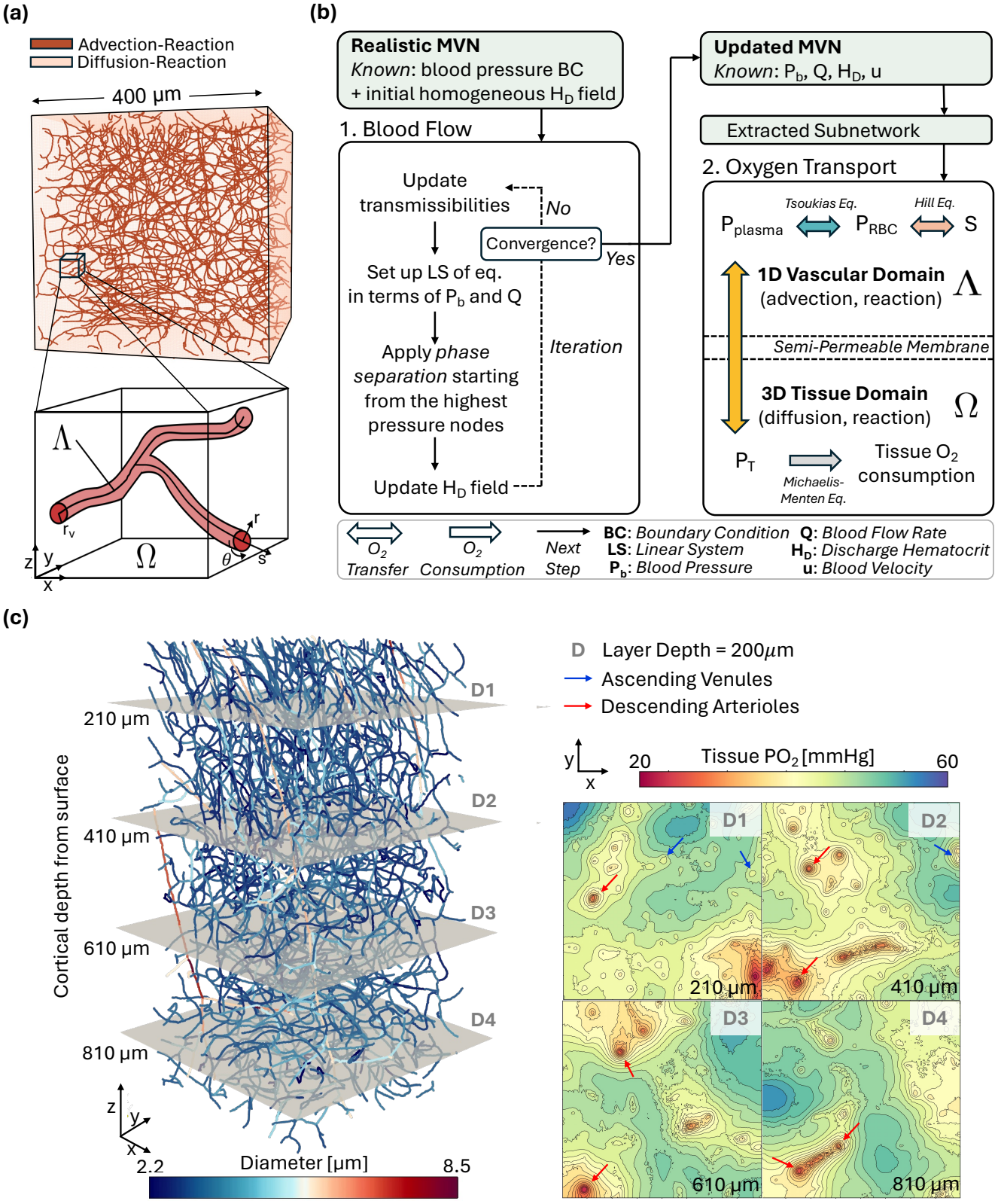
Overview of the microvascular oxygen transport modeling framework. **(a)** Schematic representation of the coupled vascular–tissue computational domain. Oxygen transport is described by advection–reaction equations in the vascular network (1D) and diffusion–reaction equations in the surrounding tissue (3D). The lower panel, illustrates a capillary bifurcation with centerline Λ embedded in the extravascular space Ω. A segment-local cylindrical coordinate system (*r, θ, s*) and a Cartesian coordinate system (*x, y, z*) are shown. The vessel radius is denoted by *r*_*v*_. **(b)** Overview of the blood flow and oxygen transport model and how they are connected. Blood flow is computed on the complete microvascular network (MVN), yielding an updated network containing blood pressure (*P*_*b*_), flow rate (*Q*), blood velocity (*v*), and discharge hematocrit (*H*_*D*_). A smaller subnetwork (top panel (a)) is then extracted from the updated network and used for oxygen transport simulations. *P*_*pl*_, *P*_*RBC*_, and *P*_*T*_, denote the oxygen partial pressure in plasma, red blood cells, and tissue, respectively, and *S* the hemoglobin oxygen saturation. **(c)** The bulk of the cortical microvascular network MVN1, spanning 400 *×*400 *×*700 *µ*m^3^. Left: Vessel segments color-coded by diameter. Right: Contour maps showing *P*_*T*_ at 4 cortical depths (D1–D4). Ascending venules and descending arterioles are highlighted in blue and red, respectively. Lower panel of (a) adapted from Koch et al. [68]

However, given that RBCs are comparable in size or often larger than the vessel diameter, they strongly affect the rheological properties of blood [59, 61] and consequently blood cannot be treated as a homogeneous Newtonian fluid in the micro-circulation. As such, three well-established phenomena must be accounted for in microvascular flow simulations: the *Fåhræus* effect, describing the reduction of tube hematocrit relative to discharge hematocrit due to the axial migration of RBCs; the *Fåhræus-Lindqvist* effect, whereby apparent blood viscosity depends on vessel diameter and hematocrit; and the *phase separation* effect, which causes unequal partitioning of RBCs and plasma at divergent bifurcations. Blood flow and hematocrit distribution are therefore mutually coupled: the apparent viscosity depends on local hematocrit and vessel diameter [62], while hematocrit is itself determined by flow partitioning at bifurcations. Since the hydraulic resistance of each vessel depends on the local apparent viscosity, the two fields cannot be determined independently, requiring an iterative solution procedure.

To compute the steady-state blood flow distribution, we employ an iterative model based on the approach of Pries et al. [63, 64] (Figure 1(b)). Rather than explicitly resolving individual RBCs, blood is modeled as a single-phase fluid, with RBCs represented through their local volume fraction, i.e., the hematocrit [27, 47, 65, 66]. The algorithm operates on realistic, experimentally derived networks, reduced to an undirected graph where vertices correspond to vessel junctions and edges to one-dimensional vessel segments [67]. The network geometry, defined by vessel lengths and diameters, together with the pressure boundary conditions at inlet and outlet nodes obtained from Schmid et al. [19], are provided as initial inputs to the model. The iterative procedure is initialized by assuming a homogeneous hematocrit distribution throughout the network, from which the initial apparent blood viscosity is determined for each vessel. Transmissibilities are then assembled, and a linear system based on mass conservation at network nodes is solved to obtain nodal pressures. Blood flow rates are subsequently computed from the resulting pressure gradients using a Poiseuille-type law. Starting from the highest-pressure nodes, hematocrit is propagated through the network and distributed at divergent bifurcations according to the empirical phase separation model of Pries et al. [63], while conserving both blood flow and RBC flux (see Online Resource 1, Section S1). The updated hematocrit distribution determines new apparent viscosities through the empirical in vitro relations of Pries et al. [62], which accounts for the Fåhræus–Lindqvist effect and in turn modifies the vessel transmissibilities.This iterative cycle is repeated until successive iterations satisfy the prescribed convergence criteria (see Online Resource 1, Section S1.1).

At convergence, the algorithm provides the steady-state distributions of pressure, blood flow, apparent viscosity, and hematocrit throughout the network. The algorithm is implemented in the open-source framework microBlooM and the resulting hemodynamic fields serve as input for the oxygen transport simulations.

### 2.2 Microvascular Networks

The two realistic microvascular networks used in this study were obtained from the vibrissa primary sensory cortex of mice [14] and employed in previous in silico studies [19, 32, 46]. MVN1 and MVN2 occupy tissue volumes of 1.6 mm^3^ and 2.2 mm^3^, respectively, and comprise approximately 12,100 and 19,300 vessel segments, classified as pial arteries (PA), descending arterioles (DA), capillaries (C), ascending venules (AV), and pial veins (PV), based on diameter and topology [19]. Capillaries account for 96% and 94% of the total vascular length in MVN1 and MVN2, respectively. The bulk of MVN1 is shown in Figure 1(c), depicting the vascular structure together with the oxygenation at different cortical depths, computed with the model described in this manuscript. The full MVNs were used to simulate blood flow, while four smaller subnet-works (400×400×400 *µ*m^3^, *∼* 1000 capillaries) were extracted from the full MVNs for the computationally expensive oxygen transport simulations (see Table S1 in Online Resource 1 for details). The smaller subnetworks were selected to reduce the computational cost and enable the serial execution of the large number of simulations required for the subsequent analyses.

### 2.3 Mixed-Dimension Embedded Model for Oxygen Transport

The oxygen model consists of two coupled domains. The extravascular compartment is represented by a three-dimensional domain Ω, corresponding to the brain parenchyma. The vascular compartment is modeled as a one-dimensional domain Λ, representing blood vessels and blood flow, where RBCs are incorporated through the hematocrit (volume fraction), while plasma constitutes the continuous fluid phase (Figure 1(a)). The two compartments are coupled through oxygen exchange across the capillary wall (transmural exchange), represented by the interface Γ. The model relies on three main assumptions: (1) axial oxygen diffusion within the vascular compartment is negligible (Pe = *uL/D≈* 42, with *u ≈*1 mm s^*−*1^, *L ≈*100 *µ*m, *D ≈*2.4 *×*10^*−*9^ m^2^ s^*−*1^), while radial diffusion is accounted for through the transmural exchange coefficients; (2) oxygen transport in both plasma and RBCs is governed by convection; and (3) oxygen consumption occurs exclusively in the tissue.

In the following sections, the models for the vascular compartment, tissue compartment, and transmural exchange are presented separately. These submodels are then integrated to obtain the full mixed-dimensional oxygen transport PDE system, solved under steady-state conditions for the oxygen partial pressures in plasma, RBCs, and tissue. This formulation is referred to as the *Single-Flux Model*. The 1D–3D oxygen transport problem is based on the blood flow and hematocrit distributions previously computed with microBlooM, as described in the previous section (Figure 1(c)).

#### 2.3.1 Oxygen Transport in the Vascular Network

Oxygen is transported in blood in three forms: dissolved in plasma, dissolved in RBCs, and bound to hemoglobin. Dissolved oxygen is described in terms of its partial pressure, 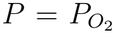 (mmHg), which is related to the oxygen concentration *C* (ml O_2_ cm^*−*3^ ) through Henry’s law [55]:

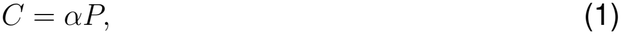

where *α* is the solubility coefficient (ml O_2_ mmHg^*−*1^ cm^*−*3^). Hemoglobin-bound oxygen is characterized by the hemoglobin saturation *S*, defined as the ratio between oxyhemoglobin and total hemoglobin concentration. Accordingly, oxygen transport in plasma, dissolved in RBCs, and bound to hemoglobin within the vascular domain Λ is described by advection–reaction equations for the cross-section-averaged variables *P*_*pl*_, *P*_*RBC*_, and *S*, respectively:

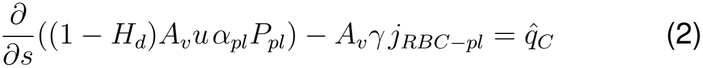

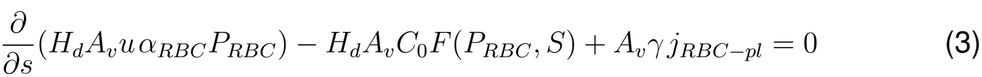

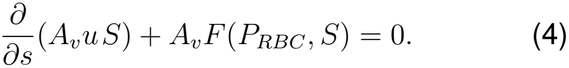

The form of Eqs. (2)–(4) follows from the balance of the total intravascular oxygen, derived in Online Resource 1, Section S3. The subscripts *pl* and *RBC* denote plasma and red blood cells, respectively. The axial blood velocity is denoted by *u*, while *H*_*d*_ represents the discharge hematocrit. Both quantities are obtained from the hemodynamic simulations (see Section 2.1) and are provided as inputs to the oxygen transport model. The variables *α, s* and *A*_*v*_ denote the oxygen solubility coefficient, the axial coordinate along the vessel centerline and the vessel cross-sectional area, respectively. The factor *γ* = 2*/r*_*v*_, where *r*_*v*_ is the vessel radius, is the vessel surface-to-volume ratio, which converts the oxygen flux per unit vessel surface into a volumetric source term along the vessel, ensuring dimensional consistency. *C*_0_ represents the oxygen binding capacity of hemoglobin, which can be expressed as 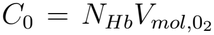, where *N*_*Hb*_ is the total heme density and 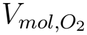 is the molar volume of oxygen at 36.9^*°*^*C*.

The term 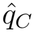 represents the oxygen exchange with the surrounding tissue and will be detailed later. The source and reaction terms accounting for oxygen exchange within the intravascular compartment, specifically *F* (*P*_*RBC*_, *S*) for hemoglobin binding and release, and *j*_*RBC−pl*_ for the oxygen flux between RBCs and plasma, are defined as shown in Eqs. (5)–(6).

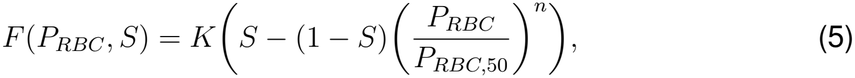

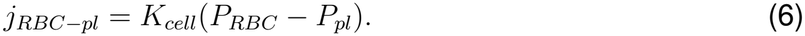

While multi-step schemes such as the Adair model [55] exist, simpler one-step approximations [26, 35, 69] are commonly adopted because reaction rates are poorly known and their additional complexity is difficult to justify. The key experimental constraint is that, at equilibrium (*F* (*P*_*RBC*_, *S*) = 0), the model must reproduce the observed hemoglobin dissociation curve, which is described by the Hill equation:

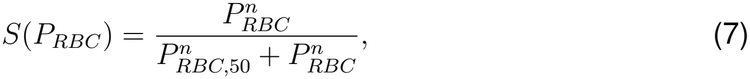

where *P*_*RBC*,50_ is the partial pressure at which hemoglobin is half-saturated, and *n* is the Hill exponent. In Eq. (5), *K* denotes the dissociation rate constant. Note that, because hemoglobin is absent in healthy plasma, the reaction term *F* (*P*_*RBC*_, *S*) is applied exclusively within the RBCs (i.e. Eqs. (3)–(4)).

On the other hand, the oxygen flux from RBCs to plasma *j*_*RBC−pl*_ is modeled using a mass transfer approach [45], in which the flux is proportional to the partial pressure difference across the RBC membrane. This representation mimics a transport mechanism in which oxygen dissolves into plasma before diffusing into the surrounding tissue. The mass transfer coefficient *K*_*cell*_ in Eq.(6) quantifies the efficiency of this exchange. A detailed analysis of *K*_*cell*_’s determinants is provided later in this work.

#### 2.3.2 Oxygen Transport Within the Tissue

Within the extravascular domain Ω, oxygen diffusion and tissue oxygen consumption are modeled by a diffusion–reaction equation governing the tissue partial pressure *P*_*T*_ :

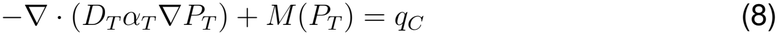

The first term represents oxygen diffusion in the tissue, where *α*_*T*_ denotes the tissue oxygen solubility and *D*_*T*_ the tissue diffusion coefficient. The source term *q*_*C*_ accounts for oxygen delivery from the vasculature to the tissue.

Cellular oxygen consumption is modeled using the Michaelis–Menten term *M* (*P*_*T*_ ) [26]:

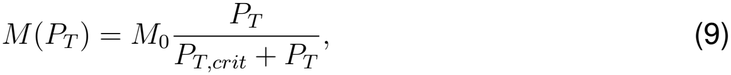

Here, *M*_0_ denotes the maximum oxygen consumption rate (*µ*m^3^ O_2_ *µ*m^*−*3^ s^*−*1^) and *P*_*T,crit*_ is the oxygen partial pressure at which the consumption rate reaches half of *M*_0_. This formulation reflects the saturable nature of oxygen consumption by tissue: at low oxygen levels, consumption increases approximately linearly with the available oxygen, while at higher levels it approaches a maximum rate, consistent with the behavior of enzymatic reactions.

#### 2.3.3 Transmural Exchange

To describe oxygen delivery from the vasculature to the surrounding tissue, the transmural flux 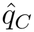 must be specified. The capillary wall, comprising the endothelial surface layer, basal membrane, and glycocalyx [70], is modeled as a semipermeable membrane and approximated as a two-dimensional interface Γ separating the vascular (Λ) and extravascular (Ω) compartments. Rather than resolving the individual layers of the wall explicitly, their resistance to oxygen transport is described as an effective transmural permeability. Consequently, the finite diffusion resistance of the wall represents the discontinuity in oxygen partial pressure across Γ. Assuming that oxygen crosses the wall by passive diffusion driven by partial pressure gradients, the transmural flux is described by the simplified Kedem–Katchalsky formulation [71]:

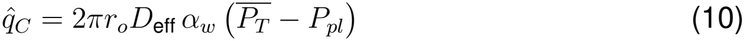

where 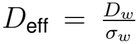 denotes the effective diffusive conductivity of the wall, which is defined as the ratio between the diffusion coefficient *D*_*w*_ and the wall thickness *σ*_*w*_. *α*_*w*_ is the oxygen solubility in the capillary wall, and *r*_*o*_ is the outer vessel radius, defined as the lumen radius plus the capillary wall thickness, *r*_*o*_ = *r*_*v*_ + *σ*_*w*_. *P*_*pl*_ is the plasma oxygen partial pressure, and 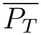 is the average tissue oxygen partial pressure at the vessel wall, defined as: 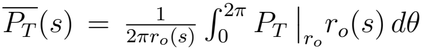, where *θ* denotes the angular coordinate in the local cylindrical coordinate system (*r, θ, z*) centered on the vessel centerline. Note that, for the vascular domain, the cross-sectionally averaged plasma partial pressure at the vessel wall coincides with the value defined on the vessel centerline in the reduced one-dimensional model [72, 73]. The transmural exchange term 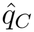 is incorporated into the three-dimensional tissue domain by reconstructing the source distribution on the lateral surface of each vessel [68]. In this formulation, the exchange term is projected onto Γ through a Dirac measure [74]. Specifically, we define

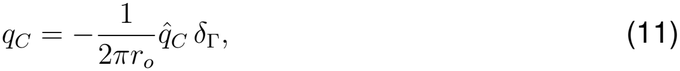

where *δ*_Γ_ is the Dirac delta distribution supported on Γ, with units of m^*−*1^, satisfying

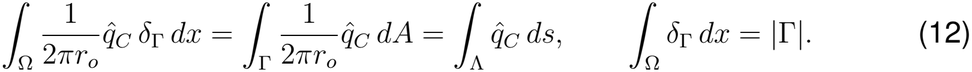

This formulation ensures mass conservation across the interface and provides a consistent approximation of the transmural exchange that would arise in a fully resolved three-dimensional description of the vessel wall.

#### 2.3.4 Initial and Boundary Conditions for Oxygen Transport

Because axial diffusion within the vessels is neglected, the vascular transport equations (Eqs. (2)–(4)) are first order in *s*, and boundary data are required only where blood enters the domain. Let *Q* be the blood flow rate through a boundary point of *∂*Λ, defined as positive when blood leaves the domain. The advective oxygen fluxes through this point are then given by

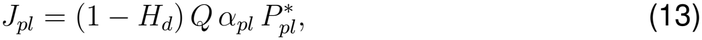

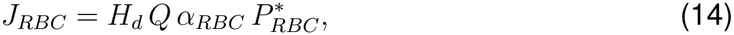

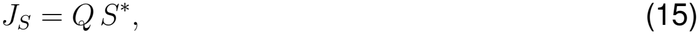

where, for *X ∈ {P*_*pl*_, *P*_*RBC*_, *S }, X*^*\**^ denotes the prescribed inflow value *X*^in^ at inflow boundaries (*Q <* 0) and the local value of *X* at outflow boundaries (*Q >* 0). Since transport is purely advective, imposing these fluxes at inflow boundaries is equivalent to prescribing 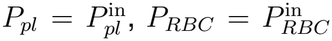 and 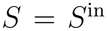, whereas at outflow boundaries oxygen is simply advected out of the domain and no condition is needed. The inflow values are assigned according to vessel type (descending arteries, ascending veins, and capillaries); for capillaries, 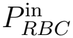 additionally depends on the discharge hematocrit (Online Resource 1, Section S4).

In the extravascular domain Ω, the outer tissue boundary *∂*Ω is subject to a homogeneous Neumann condition,

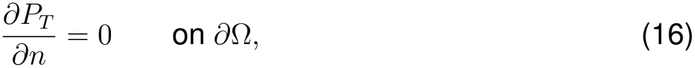

where *n* denotes the outward unit normal vector to *∂*Ω. This condition enforces zero oxygen flux across *∂*Ω, preventing exchange with the external environment and restricting tissue oxygen transport to diffusion within Ω and exchange with the embedded vascular network.

Although the problem is formulated and solved under steady-state conditions, such that the final solution is independent of the initial guess, the choice of initial conditions remains important for the numerical solution of the nonlinear system. Initial values in the vascular domain are prescribed according to the same criteria used to define the inflow boundary conditions, while the tissue oxygen partial pressure in the extravascular domain is initialized to zero.

### 2.4 Numerical Treatment and Implementation

The coupled mixed-dimensional PDE system described above is spatially discretized using a cell-centered finite volume method with a two-point flux approximation (TPFA) [75]. The resulting discrete equations are assembled into a block-matrix structure in residual form [73], yielding a nonlinear system defined by the residual function **r**(**u**), where **u** denotes the vector of primary unknowns. Since the vascular and extravascular problems are strongly coupled through the oxygen exchange terms, the discrete nonlinear system is assembled and solved monolithically using Newton’s method. The corresponding nonlinear formulation and block structure of the Jacobian are reported in Online Resource 1, Section S5.

Although the monolithic Newton method provides an efficient framework for solving the coupled nonlinear system, it does not enforce the physical constraints on the solution variables. Therefore, a projected Newton method is employed. In this approach, the primary variables are restricted to lie within prescribed bounds during the non-linear iterations. If a Newton update drives a variable outside its admissible range, it is projected back onto the closest bound. This ensures that all intermediate states remain physically meaningful, while preserving the convergence properties of the Newton method. In the present model, the hemoglobin oxygen saturation is per definition bounded in the interval

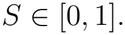

The oxygen partial pressures in plasma, red blood cells, and tissue are constrained to remain non-negative:

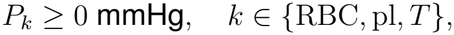

while an upper bound of 200 mmHg is imposed on *P*_RBC_ to avoid excessively large values and ensuring consistency with in vivo observations [42, 43, 76]. The need for such constraints becomes particularly evident in the low-hematocrit regime. As *H*_*d*_ *→*0, the vascular system in Eqs. (2)–(4) becomes weakly coupled and may lead to numerical illconditioning. In this limit, RBCs are almost absent and the oxygen exchange between RBCs and plasma (*j*_RBC*−*pl_) should vanish. However, *P*_RBC_ loses physical relevance, potentially taking arbitrarily large values due to the weak coupling. From a numerical standpoint, enforcing a strictly vanishing exchange flux (*j*_RBC*−*pl_ = 0) leads to a singular system matrix. As an additional regularization, the coefficient *K*_cell_ is therefore assigned a sufficiently small non-zero value (6.5 *·* 10^*−*10^ m^3^ O_2_ s^*−*1^ mmHg^*−*1^ m^*−*2^), which preserves the coupling structure of the system and avoids the singularity. Together, the box constraints and this mild regularization ensure that the oxygen exchange tends to zero as *H*_*d*_ *→*0, while guaranteeing numerical robustness and physical consistency over the full range of hematocrit values. Regardless of these numerical treatments, *P*_RBC_ and *S* are not defined in segments with exactly *H*_*d*_ = 0, where no RBCs are present, and their values there are excluded from all statistics and figures reported in this work.

The model is implemented in the open-source software framework DuMux [75], which is based on the Distributed Unified Numerics Environment (DUNE) [77]. The mixed-dimension embedded oxygen transport model builds upon a DuMux application for multidomain porous media problems [74].

Grid convergence was verified by comparison with a reference solution computed on a very fine grid (see Online Resource 1, Section S6). Based on the results of the convergence study, the 3D computational grid was discretized with a cell size of *µ*m. At this resolution, the runtime for a single forward simulation is approximately 20 minutes on a standard laptop (see Online Resource 1, Section S6).

### 2.5 Mass Transfer Coefficient for RBC–Plasma Oxygen Exchange

To assess the influence of the RBC–plasma MTC on oxygen transport and tissue oxygenation, we consider analytical expressions for *K*_cell_ derived from previous computational studies that explicitly resolve individual RBCs, enabling detailed characterization of microscale oxygen exchange [35, 37]. The formulation proposed by Lücker et al. [35] characterizes the dependence of the MTC on hematocrit, whereas the model of Vadapalli et al. [37] accounts for dependencies on both hematocrit and oxygen saturation.

Within the range *H*_*t*_ *∈* [0.07, 0.50], with *H*_*t*_ denoting the tube hematocrit, the data extracted from Lücker et al. [35] were fitted using a fifth-degree polynomial:

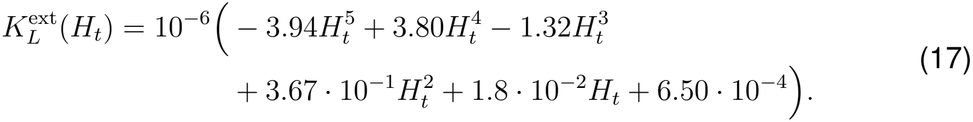

For hematocrit values below the fitted range, i.e. *H*_*t*_ *∈* [0, 0.07), the polynomial decreases monotonically towards 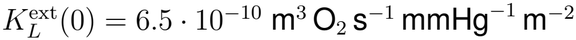.

For hematocrit values beyond the available data range, i.e. *H*_*t*_ *>* 0.5, the function is extended with the tangent to Eq. (17) at *H*_*t*_ = 0.50:

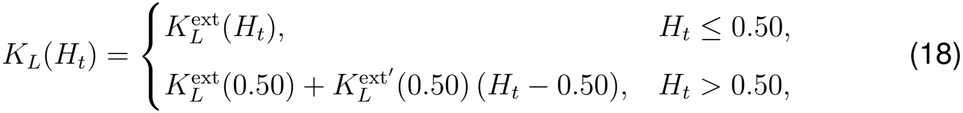

where 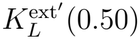 denotes the derivative of Eq. (17), evaluated at *H*_*t*_ = 0.50. The resulting piecewise-defined function *K*_*L*_(*H*_*t*_) is illustrated in Figure 2(a).

**Fig. 2.**
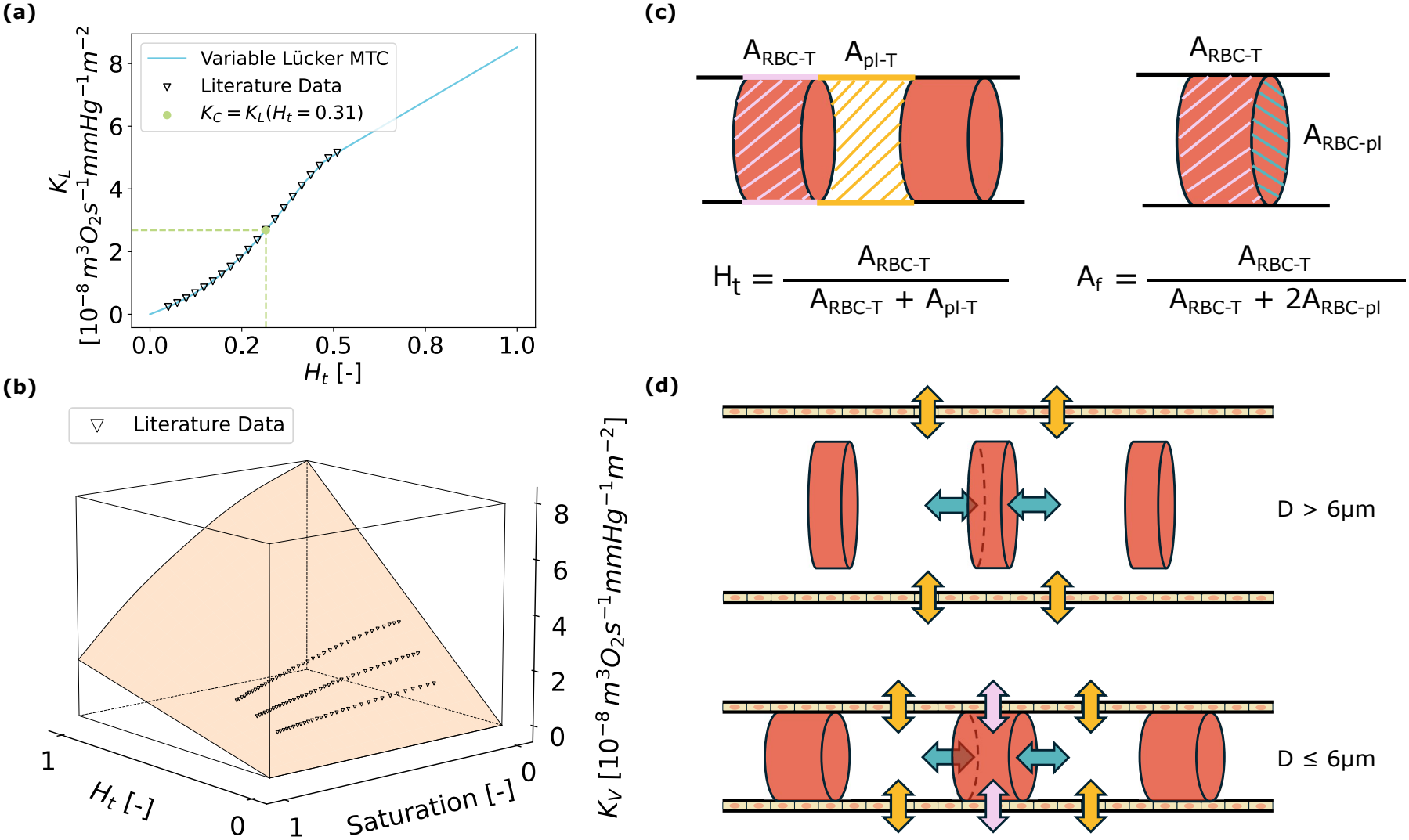
Oxygen exchange pathways in the single-flux and double-flux models and related mass transfer coefficients. **(a)** Hematocrit-dependent mass transfer coefficient *K*_*L*_, fitted to literature data [35] for *H*_*t*_ *∈* [0.07, 0.5] and extended linearly for *H*_*t*_ *>* 0.5 (see Eq. (18)). **(b)** Hematocrit-and saturation-dependent mass transfer coefficient *K*_*V*_, obtained by fitting the data reported by Vadapalli et al. [37] (*S ∈* [0.10, 0.80] at *H*_*t*_ = 0.20, 0.30, 0.40) and extrapolated over *H*_*t*_, *S ∈* [0, 1]. **(c)** Partitioning of the exchange surfaces associated with the direct RBC–tissue pathway. The schematic illustrates the surface fractions assigned to the different transport pathways and the corresponding scaling factors (*H*_*t*_ and *A*_*f*_ ) used to rescale the oxygen fluxes according to the partitioned exchange areas. **(d)** Schematic representation of the oxygen fluxes considered in the double-flux model. For vessels with diameter larger than 6 *µ*m (top), oxygen exchange occurs through the conventional RBC–plasma (green arrow) and plasma–tissue (yellow arrow) pathways only. For vessels with diameters smaller than 6 *µ*m (bottom), a direct RBC–tissue oxygen flux (pink arrow) is additionally activated

The MTC based on data from Vadapalli et al. [37] was extracted as a function of oxygen saturation *S ∈* [0.10, 0.80] for three discrete values of tube hematocrit, namely *H*_*t*_ = 0.20, 0.30, and 0.40. The extracted data were fitted using the following expression:

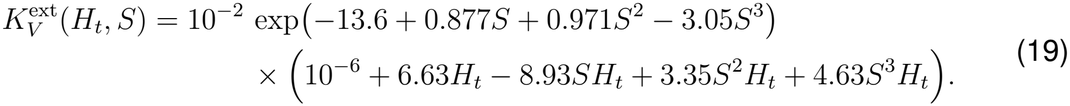

This function ensures non-negative values and captures the trends observed in the data, exhibiting a monotonic increase with respect to *H*_*t*_ for fixed values of *S*, and a nonlinear decrease with respect to *S* for fixed *H*_*t*_. For small hematocrit values, Eq. (19) approaches zero. To ensure numerical robustness, a smooth lower bound was imposed,

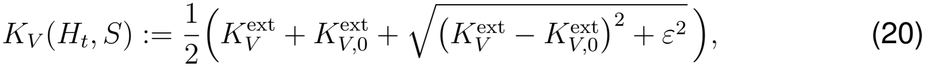

With

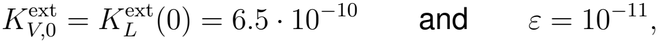

both expressed in m^3^ O_2_ s^*−*1^ mmHg^*−*1^ m^*−*2^. The resulting function *K*_*V*_ (*H*_*t*_, *S*) is shown in Figure 2(b), illustrating its variation with respect to both tube hematocrit and oxygen saturation over the full domain *H*_*t*_, *S ∈* [0, 1] (representative cross-sections of the reconstructed surface at three discharge hematocrits are also available in Figure 3(b)).

In the subsequent analyses, three RBC–plasma mass transfer coefficient formulations are compared: constant, hematocrit-dependent [35], and hematocrit–saturation dependent [37]. The first approach assumes a constant coefficient,

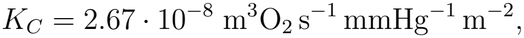

applied uniformly throughout the domain (corresponding to *K*_*L*_ evaluated at the mean network hematocrit, *H*_*t*_ = 0.31). The conversion from tube hematocrit (*H*_*t*_), used in the original MTC formulations, to discharge hematocrit (*H*_*d*_), used in the present model, is performed according to the empirical relationship proposed by Pries et al. [63].

**Fig. 3.**
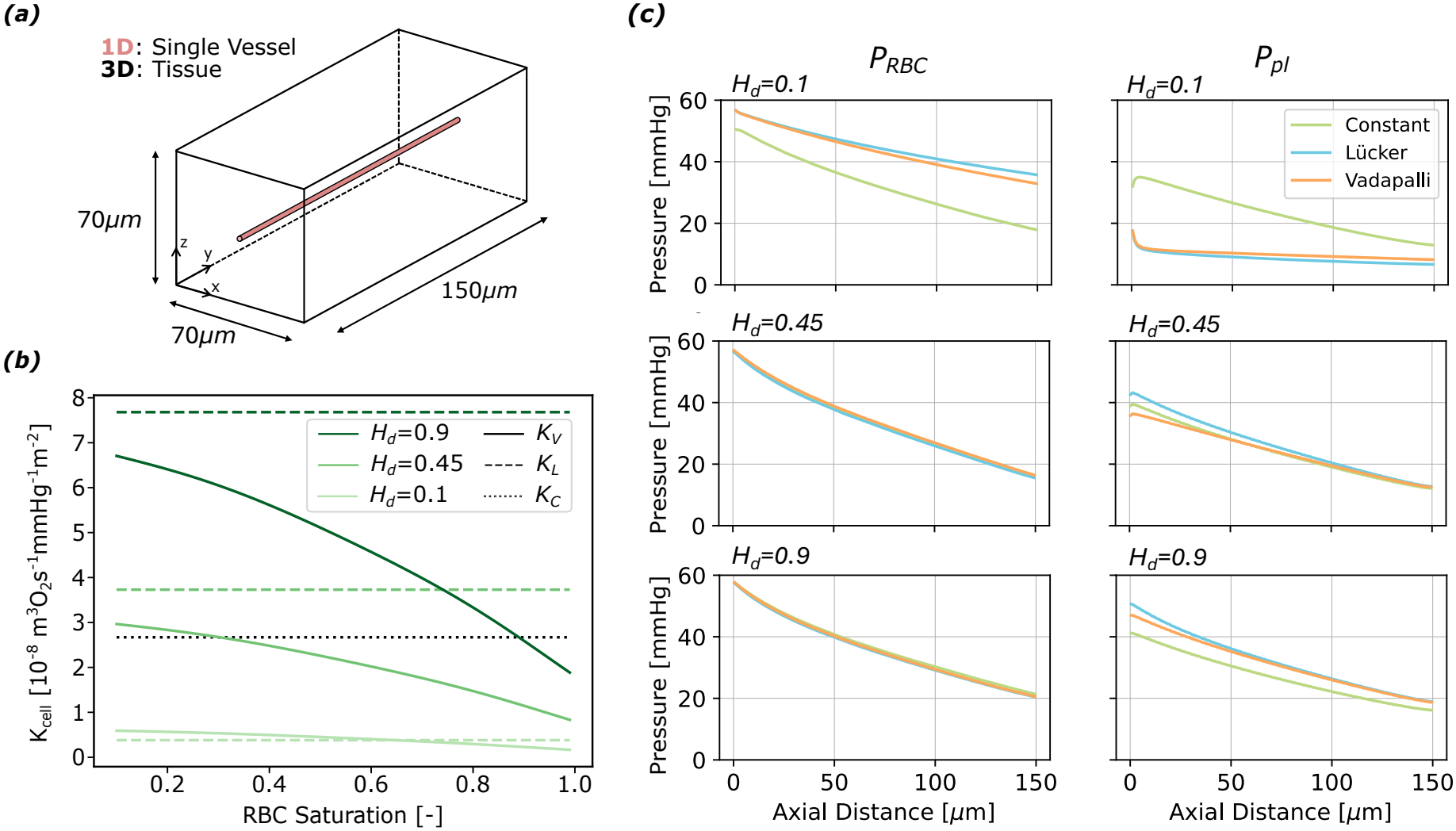
Influence of different mass transfer coefficient formulations on oxygen partial pressure in single-vessel simulations. **(a)** Computational setup of the single-vessel simulations. **(b)** Mass transfer coefficients reconstructed from the functions *K*_*V*_ (solid lines) and *K*_*L*_ (dashed lines) fitted to the data reported by Vadapalli et al. [37] and Lücker et al. [35], respectively. The fitted coefficients are shown as a function of oxygen saturation for three different discharge haematocrit values (*H*_*d*_ = 0.1, 0.45, and 0.9). The dotted line denotes *K*_*C*_, corresponding to *K*_*L*_(*H*_*t*_ = 0.31). **(c)** Profiles of red blood cell oxygen partial pressure (*P*_RBC_, left column) and plasma oxygen partial pressure (*P*_pl_, right column) as a function of the axial distance along the vessel centerline. Results for three different MTC formulations are compared: *Constant* (green), *Lücker* (light blue), and *Vadapalli* (orange). Each row corresponds to a different discharge haematocrit: *H*_*d*_ = 0.1 (top row), *H*_*d*_ = 0.45 (middle row), and *H*_*d*_ = 0.9 (bottom row)

### 2.6 Extension to a Double-Flux Model: Inclusion of Direct RBC–Tissue Exchange

In the *Double-Flux Model*, in addition to the classical two-step pathway (RBC *→*plasma *→*tissue), we consider a direct oxygen flux from RBCs to the surrounding tissue. This additional pathway is also driven by partial pressure gradients but is modulated by the combined resistance of the RBC membrane and the capillary wall.

In the baseline model, the entire capillary surface is available for plasma–tissue exchange (Eq. 10). With the introduction of the RBC–tissue pathway, the capillary surface is partitioned into two exchange regions associated with plasma–tissue and RBC–tissue oxygen transfer, respectively. The fraction of the exchange surface assigned to each pathway is assumed to follow the tube hematocrit *H*_*t*_, defined as the ratio between the RBC volume and the total blood volume within the vessel. Accordingly, *H*_*t*_ is used as a proxy for the fraction of the capillary surface associated with RBC-mediated exchange:

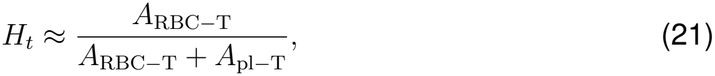

where *A*_RBC*−*T_ and *A*_pl*−*T_ denote the effective capillary surface areas assigned to RBC–tissue and plasma–tissue exchange, respectively (Figure 2(c)). Consequently, a fraction *H*_*t*_ of the capillary surface is assigned to RBC–tissue exchange, while the remaining fraction (1 *− H*_*t*_) is assigned to plasma–tissue exchange. The corresponding source terms are therefore scaled by *H*_*t*_ and (1 *− H*_*t*_), respectively.

In the single-flux formulation, the entire RBC surface is assumed to contribute to RBC–plasma exchange. In the double-flux model, instead, red blood cells are allowed to exchange oxygen simultaneously with both plasma and tissue. To represent this mechanism, each RBC is idealized as a cylinder of constant volume *V*_RBC_, whose geometry depends on the local vessel radius *r*_*v*_. Under this assumption, the RBC surface is partitioned into two contributions: *A*_RBC*−*T_, associated with RBC–tissue exchange, and the two circular end faces *A*_*RBC−pl*_, associated with RBC–plasma exchange. The corresponding surface areas are given by

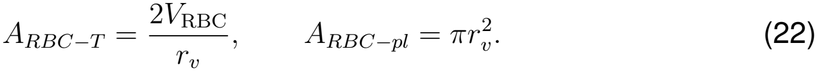

The fraction of RBC surface involved in RBC–tissue exchange is then defined as

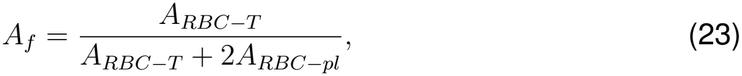

while the remaining fraction (1*− A*_*f*_ ) contributes to RBC–plasma exchange (Figure 2(c)).

The direct RBC–tissue oxygen flux is assumed to be proportional to the partial pressure difference between RBCs and tissue. The corresponding proportionality coefficient is derived using a resistance-in-series formulation based on the Meyer–Overton law [78], accounting for transport across both the RBC membrane and the capillary wall:

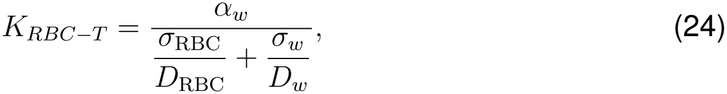

where *σ* denote membrane thicknesses, and *D* the respective diffusion coefficients. According to the expression in Eq. 10, the corresponding source term for the RBC– tissue oxygen flux is defined as:

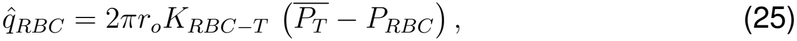

and distributed in the tissue domain as:

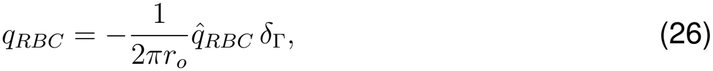

where *δ*_Γ_ is defined analogously to Eq. 12.

The direct RBC–tissue flux is incorporated only in vessels with diameters *≤* 6 *µ*m (Figure 2(d)), corresponding to the regime in which vessel dimensions become comparable to the characteristic size of RBCs [61]. In this range, the reduced separation between RBCs and the vessel wall motivates the investigation of an additional direct oxygen exchange pathway. Accordingly, the inclusion of this mechanism modifies the governing equations only for vessels where RBC–tissue exchange is active, whereas for larger vessels the baseline formulation remains unchanged (Eqs. (2)–(12)). Under these conditions, the modified governing system is formulated as follows:

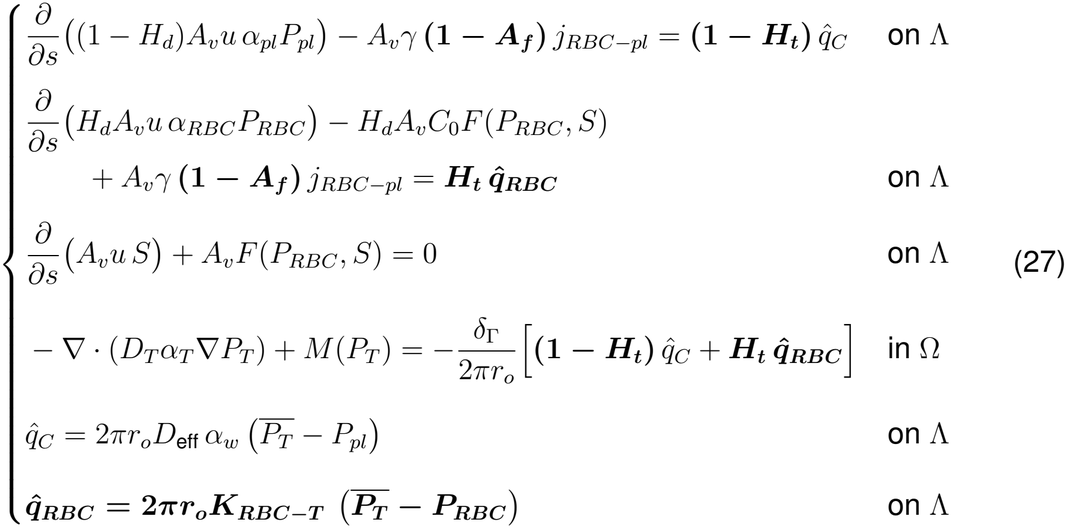

Terms in bold are introduced by the double-flux model, while all other terms are taken unchanged from the single-flux formulation (Eqs. (2)–(12)).

## 3 Results

All results presented in the following have been obtained with the parameter values reported in Table S4 (Online Resource 1), unless specified otherwise within the corresponding sections.

### 3.1 Single-Flux Model

#### 3.1.1 Variable MTCs are most relevant at low hematocrit

To isolate the impact of the different mass transfer coefficients, single-vessel simulations were performed. The computational domain consists of a 150 *µ*m straight vessel with a fixed diameter of 4 *µ*m embedded centrally in a 70 *×* 70 *×* 150 *µ*m^3^ tissue domain (Figure 3(a)). The domain size was selected to be consistent with typical inter-capillary spacing in cortical tissue and characteristic oxygen diffusion distances in consuming tissue [27, 79]. The tissue domain is discretized using a Cartesian grid with a spatial resolution of 2.5 *µ*m, while the vascular domain is discretized with a resolution of *∼*1 *µ*m. For the single-vessel configuration, inflow boundary conditions are prescribed using fixed values consistent with physiological capillary conditions, as summarized in Online Resource 1, Table S5. The discharge hematocrit *H*_*d*_ is constant along the vessel, and the blood flow rate is adjusted for each hematocrit level so that the total oxygen inflow is the same in all simulations and slightly exceeds the maximal tissue oxygen demand (Online Resource 1, Table S5), preventing differences in oxygen supply from affecting the comparison. Three hematocrit levels are tested to comment on the impact of the different MTC formulations. The corresponding *K*_*cell*_ values for Lücker and Vadapalli are shown in Figure 3(b). Single-vessel simulations revealed that, at low hematocrit (Figure 3(c), first row), the MTC formulation strongly affects the oxygen profiles. With the Lücker and Vadapalli formulations, *P*_*RBC*_ decreases moderately along the vessel, from *∼*59 mmHg to 35 mmHg, with slightly lower values for Vadapalli. By contrast, the constant MTC formulation produces a substantially steeper decline, with *P*_*RBC*_ dropping to about 18 mmHg over the same distance. Correspondingly, the constant MTC predicts higher *P*_*pl*_ values along the entire vessel, whereas the variable MTCs yield lower but more uniform plasma oxygenation. This indicates that, at low hematocrit, the variable MTCs slow down the release of oxygen from the RBCs, distributing it more uniformly along the vessel. For intermediate and high hematocrit values (Figure 3(c), second and third row), the influence of the MTC formulation on *P*_*RBC*_ is reduced, with nearly indistinguishable profiles for the three formulations. Also for *P*_*pl*_ the variations between formulations are smaller than under low hematocrit conditions. This reduced sensitivity to the MTC formulation at high hematocrit can be understood by considering that oxygen released by the RBCs must overcome two resistances in series: the RBC–plasma interface, characterized by the MTC, and the capillary wall. Since the oxygen inflow is the same in all simulations, hematocrit does not alter the oxygen supply, but determines the number of RBCs per unit vessel length and thus the interfacial area available for exchange. At low *H*_*d*_, this area is small and the RBC–plasma interface is the dominant resistance, so that the MTC controls oxygen release. As *H*_*d*_ increases, the interfacial resistance decreases until the capillary wall becomes the limiting step, making the oxygen profiles largely insensitive to the choice of MTC even though its values still differ considerably.

#### 3.1.2 Similar global oxygenation masks differences in hypoxic burden between MTC formulations

We now extend the analysis to realistic microvascular networks, where the interplay between heterogeneous flow distribution, vascular geometry, and oxygen transport mechanisms introduces additional variability compared to the single-vessel configuration. For this analysis, four representative subnetworks were considered, with two extracted from MVN1 and two from MVN2 (Online Resource 1, Table S1). The tissue domain is discretized using a uniform structured grid of 160 *×*160*×* 160 elements, corresponding to a spatial resolution of 2.5 *µ*m.

The analysis focuses on quantifying whether different MTC formulations translate into measurable variations in global oxygen availability. The frequency histograms of *P*_*T*_ (Fig. 4(a)) show that the overall oxygenation profiles are comparable across the three MTC formulations, with two peaks around *∼*25 mmHg and *∼*40 mmHg, reflecting the different oxygenation levels of the subnetworks. The mean *P*_*T*_ for all formulations is between 31 and 34 mmHg, consistent with previously reported experimental observations [21]. This overall agreement indicates that the choice of MTC formulation has a limited impact on the global tissue oxygenation.

**Fig. 4.**
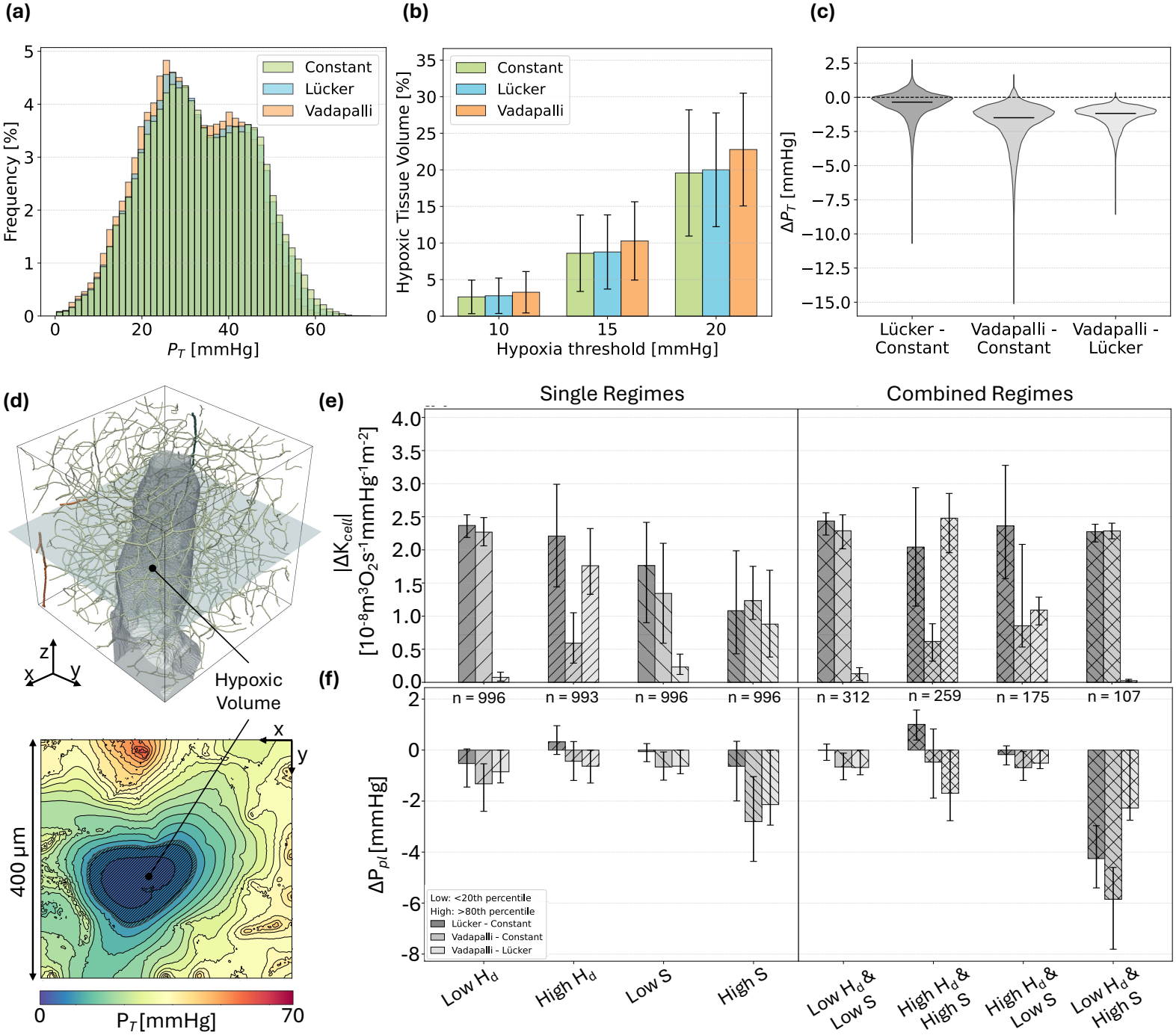
Impact of RBC–plasma mass transfer coefficient formulations on tissue oxygenation and hypoxic burden in realistic microvascular networks. **(a)** Frequency distributions of *P*_T_ predicted by the three MTC formulations. **(b)** Fraction of hypoxic tissue volume predicted by the three MTC formulations for hypoxia thresholds of 10, 15, and 20 mmHg. Only three subnetworks exhibiting non-zero hypoxic tissue volume are shown; the remaining one showed 0% hypoxic tissue volume for all thresholds and all MTC formulations. **(c)** Violin plots of pairwise differences in tissue oxygen partial pressure, Δ*P*_T_, obtained from the comparisons Lücker–Constant, Vadapalli– Constant, and Vadapalli–Lücker. **(d)** Visualization of oxygenation patterns and hypoxic regions in representative realistic microvascular network. The vascular network highlights the different vessel types (arteries, orange; veins, blue; capillaries, light green), while the corresponding *P*_T_ (obtained with Vadapalli’s formulation) is shown on a representative cross-sectional plane (bottom). The hatched region indicates hypoxic tissue defined by *P*_T_ *<* 20 mmHg. **(e)** Absolute differences in the mass transfer coefficient, |Δ*K*|, obtained from pairwise comparisons of simulations with the three different MTC formulations. Results are grouped according to low/high discharge hematocrit (*H*_*d*_) and oxygen saturation (*S*) regimes (left), where low refers to values *<*20^th^ percentile and high to values *>*80^th^ percentile. On the right, different combinations of those regimes are shown. **(f)** Difference in plasma oxygen partial pressure, Δ*P*_pl_, for the same pairwise comparisons and local flow conditions as shown in panel (e). Bar plots and error bars represent mean and standard deviation across all vessels of the respective group. Results shown in panels (a), (c), (e), and (f) were obtained by pooling data across the four realistic microvascular subnetworks

However, the overall similarity in the *P*_*T*_ distribution hides that relevant differences exist if the focus is shifted to metrics like tissue hypoxia. Both variable MTC models consistently predict higher hypoxic fractions compared to the constant case across all hypoxic thresholds (Figure 4(b)). While the Lücker model yields values very close to the constant case, the Vadapalli model exhibits the highest hypoxic fractions, with differences that become more pronounced as the threshold increases. Figure 4(d) shows an example of the resulting oxygenation pattern and hypoxic regions for a single MVN, obtained using Vadapalli’s formulation and a hypoxia threshold of *P*_*T*_ *<* 20 mmHg. This indicates that even relatively small shifts in *P*_*T*_ can translate into significant differences in hypoxic burden, not captured by global distributions alone.

This is further confirmed by the pairwise variations in *P*_*T*_ (Fig. 4(c)). Comparing the variable MTC formulations with the constant model shows that the distributions are shifted toward negative values, meaning that the constant MTC generally predicts higher oxygen levels. The small shift of the Lücker–Constant distribution is expected, since the constant MTC corresponds to the Lücker formulation evaluated at the subnetworks’ mean hematocrit. Noteworthy are the pronounced negative peaks of about *−* 11 mmHg and *−* 15 mmHg for the Lücker–Constant and Vadapalli–Constant comparisons, highlighting the strong local impact of the MTC formulation. The comparison between the Vadapalli and Lücker formulations yields a narrower distribution, still predominantly shifted toward negative values.

Overall, these results highlight the relevance of correctly assigned MTCs for accurate predictions of the local oxygenation and hypoxic fraction, which remain hidden in global partial pressure distributions.

#### 3.1.3 Differences between MTC formulations in realistic microvascular networks emerge in specific hematocrit and saturation regimes

The previous analysis showed that different MTC formulations can lead to systematic variations in predicted oxygen levels, despite exhibiting similar global distributions. To identify the physiological conditions responsible for these discrepancies, we now examine their dependence on local hematocrit and oxygen saturation levels.

The comparison of *K*_*cell*_ between the Constant and Lücker formulations shows large differences at both low and high hematocrit (Figure 4(e), see also Figure 2(a)). Oxygen saturation has a notable but smaller influence. The effect on the oxygen field, however, is not equivalent to the behavior of the MTC (Figure 4(f)). Despite the large differences in *K*_*cell*_, the Lücker–Constant variations in *P*_*pl*_ remain below about 1 mmHg in most categories. The most pronounced effect occurs for the combination of low hematocrit and high saturation, with *P*_*pl*_ differences of about 4 mmHg, consistent with the findings discussed in the previous section.

The comparison between the Constant and Vadapalli formulations shows a strong dependence on hematocrit. The largest differences between *K*_*V*_ and *K*_*C*_ are observed at low hematocrit, consistently across saturation levels, while at high hematocrit the Vadapalli MTC remains close to the constant value. This is reflected in the Δ*P*_*pl*_ distributions, where the differences are larger in low-*H*_*d*_ than in high-*H*_*d*_ regions and are further amplified in high-saturation regimes, reaching their maximum when low hematocrit and high saturation are combined. Overall, the constant MTC formulation tends to overestimate oxygen availability (up to *∼*6 mmHg), particularly in low-hematocrit and highly saturated regions.

The comparison between the Lücker and Vadapalli formulations highlights the role of oxygen saturation. At low hematocrit, the two formulations produce very similar coefficients across all saturation levels, indicating a weak dependence on *S*. At high hematocrit, however, the Vadapalli model introduces a stronger saturation dependence, and the difference from the Lücker MTC becomes largest at high saturation. Despite this, the resulting variations in *P*_*pl*_ remain moderate (up to about 2 mmHg), with the largest shifts observed in high-saturation regimes.

Overall, the discrepancies between the MTC formulations are not uniformly distributed. Although the coefficients differ substantially in most categories, the resulting differences in *P*_*pl*_ concentrate in high-saturation regions, particularly when combined with low hematocrit, where the constant MTC overestimates oxygen availability by several mmHg. In the remaining regions, the differences between formulations are generally within about 1 mmHg, leading to comparable predictions of oxygen partial pressure.

#### 3.1.4 The impact of MTC formulations is tied to low hematocrit regions along vascular paths

To investigate how MTC formulations influence oxygen transport along individual vascular paths, three representative flow paths were selected within one microvascular subnetwork (Fig. 5(a)), each capturing a distinct hematocrit regime. The corresponding hematocrit profiles are shown in Figure 5(b) and include a predominantly low-hematocrit path (left), a path exhibiting a transition from high to low hematocrit values (center), and a predominantly high-hematocrit path (right).

**Fig. 5.**
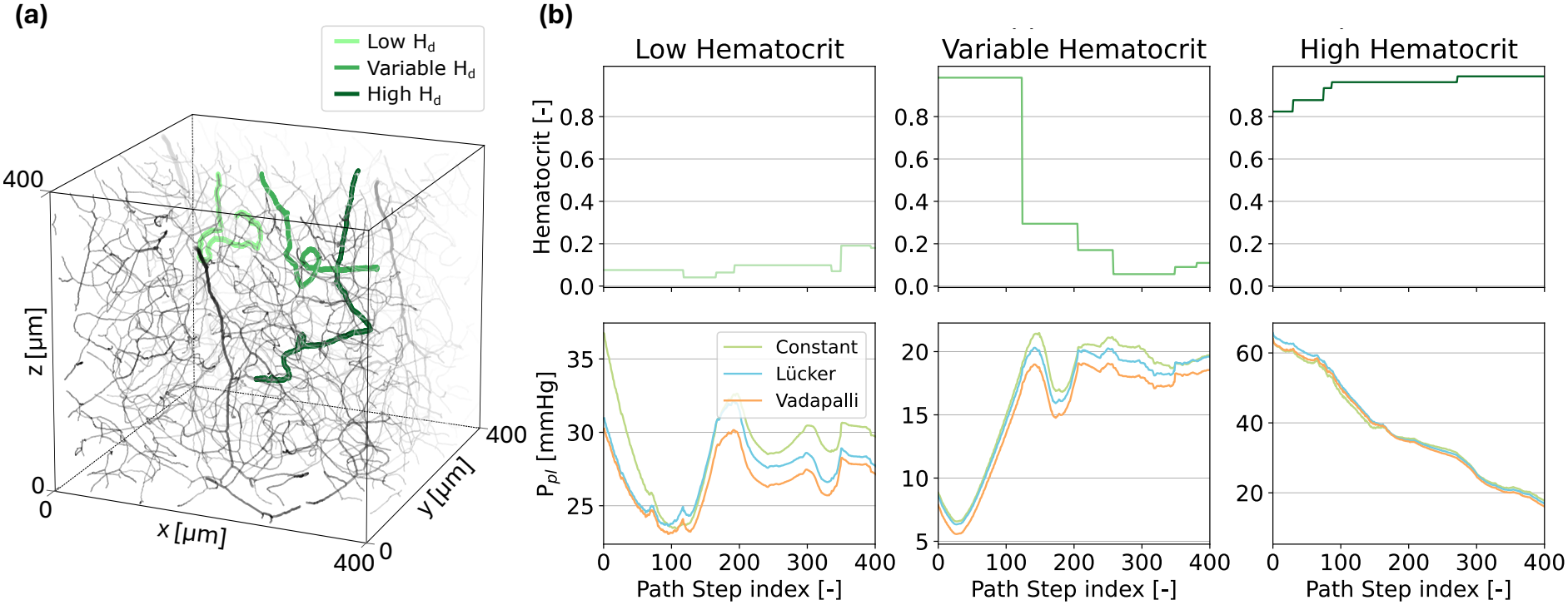
Impact of different MTC formulations along representative microvascular flow paths. **(a)** Visualization of three selected flow paths within the selected realistic microvascular network. The highlighted paths correspond to the low-, variable-, and high-hematocrit cases. **(b)** Hematocrit (top) and *P*_pl_ (bottom) profiles along three representative flow paths within the microvascular network, depicted in function of the path step index. The path step index denotes the sequential position of each 1D discretization cell along the selected flow path. For each path, results obtained using three different mass transfer coefficient formulations are compared

For the low hematocrit path the discharge hematocrit remains below *H*_*d*_ = 0.20 with an average value of *H*_*d*_ = 0.09. Consistent with the results obtained in the singlevessel configuration, the choice of MTC formulation has a noticeable impact under low-hematocrit conditions. In particular, the Constant model predicts systematically higher values of *P*_pl_ than the two variable-MTC formulations, with differences reaching up to 6 mmHg. For the high hematocrit path the discharge hematocrit remains above *H*_*d*_ = 0.80 along the entire path, with an average value of *H*_*d*_ = 0.96 (Fig. 5(b)). Despite the pronounced differences in the magnitude of the MTCs, all three formulations yield nearly identical oxygen partial pressure distributions. The third selected path exhibits substantial hematocrit variability along its length, capturing the typical heterogeneity of microvascular flow. The hematocrit remains high during the first part of the path and then decreases markedly around step 125. Beyond this point, *H*_*d*_ stays predominantly below the network-average value of 0.37. The largest differences between the three formulations are observed in the low-hematocrit region, where the Constant model yields noticeably different oxygen partial pressures. In the initial high-hematocrit segment, by contrast, the discrepancies are smaller, and the three models yield more similar predictions. This observation confirms that, under high-hematocrit conditions, oxygen transport becomes largely insensitive to the specific choice of RBC–plasma mass transfer coefficient, in agreement with the trends observed in the single-vessel simulations. Additional results showing the corresponding *P*_RBC_ profiles and the variations of the MTC coefficients (*K*_*C*_, *K*_*L*_, and *K*_*V*_ ) along the selected vascular paths are provided in Online Resource 1, Fig. S2.

### 3.2 Double-Flux Model

#### 3.2.1 Contribution of the RBC–tissue pathway to tissue oxygenation

We now use the single-flux model as reference case that is compared against the double-flux formulation, in which the RBC–tissue exchange is activated. The RBC– plasma exchange is described using the constant MTC (*K*_C_) in both models, so that differences reflect only the contribution of the additional RBC–tissue transport pathway. The *D*_RBC_ used to define the RBC–Tissue MTC was derived from *K*_C_ by invoking the Meyer–Overton law [78] and accounting for the oxygen solubility in plasma and the effective thickness of the RBC membrane (see Online Resource 1, Table S4). This choice ensures consistency between the RBC–plasma and RBC–tissue transport pathways, as both MTCs are then based on the same underlying RBC diffusivity. In the single-vessel configuration (introduced in the previous section), two parametric studies are considered: (i) fixed hematocrit (*H*_*t*_ = 0.31, corresponding to the average value observed in capillaries of realistic MVNs) with varying vessel diameter, and (ii) fixed diameter (*D* = 4 *µ*m) with varying hematocrit. As in the previous section, the blood flow rate is adjusted to the hematocrit so that the total oxygen inflow is the same in all simulations. For *D* 6 *≤ µ*m, i.e. where the second flux is active, both the vessel diameter and hematocrit affect the effective exchange area through the surface partitioning factor *A*_*f*_ (Eq. 23) and the capillary surface availability *H*_*t*_ (Eq. 21).

For vessels with 5 *µ*m *≤D ≤*6 *µ*m, activation of the direct RBC–tissue pathway leads to a moderate increase in tissue oxygen partial pressure and a negligible decrease in RBC oxygen partial pressure (Figure 6(a), top row). The trend is reversed for *D ≤* 4 *µ*m, where oxygen is retained to a greater extent within the RBCs and delivered less efficiently to the tissue. Concretely, *P*_*T*_ decreases by about 9–11% at *D* = 4 *µ*m and by 30–34% at *D* = 3 *µ*m, while *P*_RBC_ increases by up to 0.8% and 8%, respectively. For both diameters, the deviations grow along the capillary, owing to the cumulative effect of the reduced oxygen release. In all cases, the effect on *P*_*T*_ exceeds in magnitude that on *P*_RBC_, reflecting the greater sensitivity of the tissue compartment to changes in the exchange configuration. The opposite trends for 5 *µ*m*≤ D ≤*6 *µ*m and *D ≤*4 *µ*m, result from the competition between two effects of the double-flux formulation. On the one hand, the surface partitioning (Figure 6(b)) reduces the efficiency of the RBC–plasma–tissue pathway, by restricting RBC–plasma exchange to the fraction (1 *−A*_*f*_ ) of the RBC surface and plasma–tissue exchange to the fraction (1 *−H*_*t*_) of the capillary wall. On the other hand, over the remaining fraction *H*_*t*_ of the wall, oxygen is transferred directly from the RBCs to the tissue. This pathway carries a lower MTC (Eq. (24)), as it must overcome the resistance of both the RBC membrane and the capillary wall, but is driven by a larger partial pressure difference, since *P*_RBC_ *> P*_*pl*_. The balance between the two effects depends strongly on vessel diameter through *A*_*f*_ (Eq. (23)), which increases as *D* decreases. For 5 *≤ µ*m *D ≤* 6 *µ*m, the residual RBC–plasma exchange surface still ensures an adequate oxygen supply to the plasma, and the direct pathway more than compensates for the reduced plasma– tissue exchange area. For *D ≤* 4 *µ*m, RBC–plasma exchange becomes so restricted that it limits the oxygen supply to the plasma, impairing the plasma–tissue pathway. The direct pathway, confined to the fraction *H*_*t*_ of the wall and limited by its lower MTC, can no longer compensate for this loss. The nonlinear dependence on diameter highlights the complex interplay between the two exchange pathways and their combined effect on oxygen discharge.

**Fig. 6.**
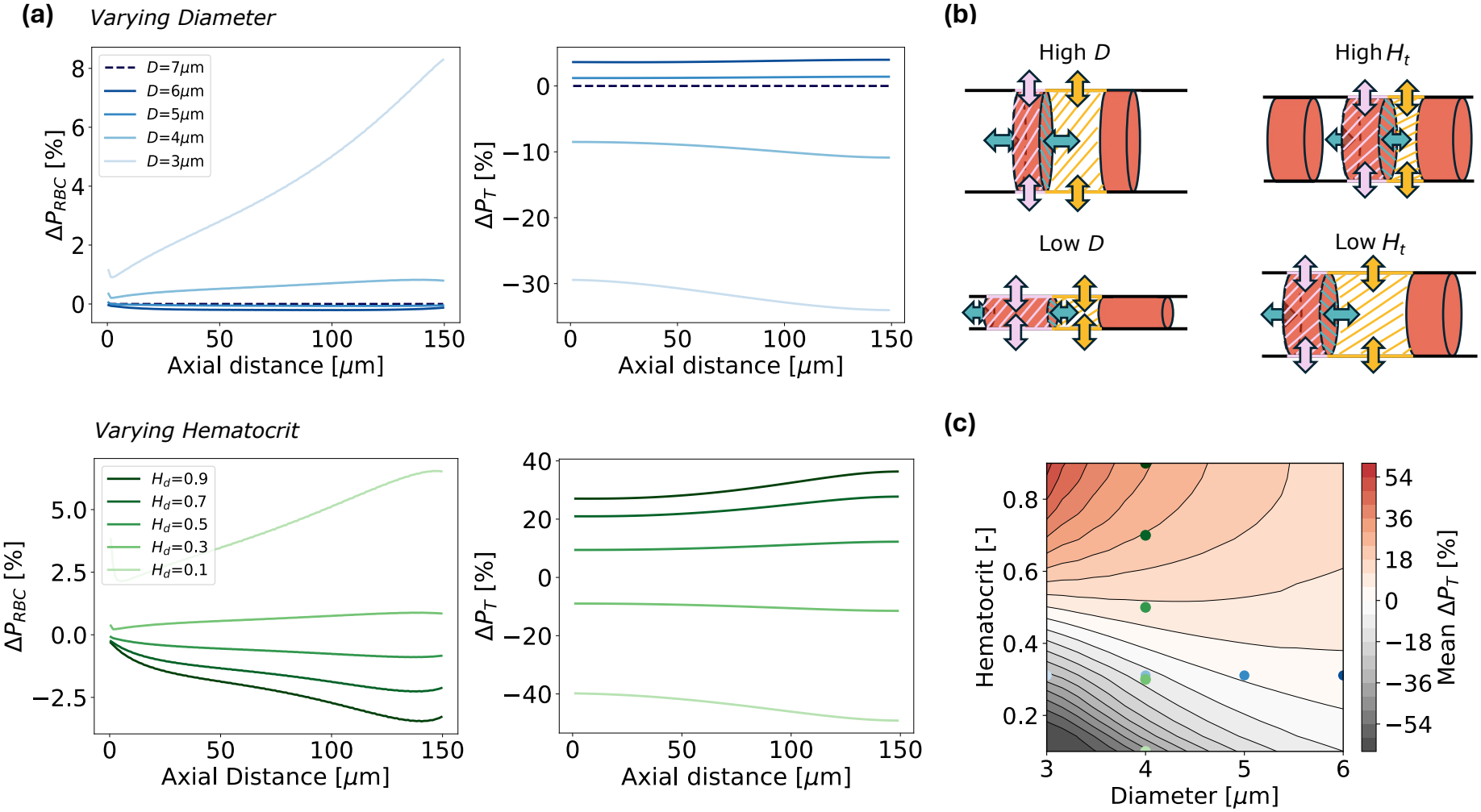
Effects of the double-flux model on oxygen partial pressure distributions in single-vessel simulations. **(a)** Relative variation in red blood cell oxygen partial pressure (*P*_RBC_; left column) and tissue oxygen partial pressure (*P*_T_; right column) as a function of the axial distance along the vessel centerline. The top row presents the results obtained by varying the vessel diameter while keeping the hematocrit fixed at *H*_*t*_ = 0.31. The bottom row shows the results obtained by varying the discharge hematocrit while maintaining a constant vessel diameter of 4 *µ*m.**(b)** Schematic representation of the oxygen fluxes considered in the double-flux model and the corresponding partitioning of the exchange surfaces. Four configurations are shown: low diameter (Low D), high diameter (High D), low hematocrit (Low *H*_*t*_), and high hematocrit (High *H*_*t*_). **(c)** Contour map of the mean percentage variation in *P*_T_ predicted by the double-flux model relative to the single-flux model, in function of vessel diameter and hematocrit. All variations are computed relative to the corresponding results obtained with the original single-flux model. Colored markers indicate the parameter combinations explored in panel (a)

Hematocrit modulates the double-flux response through the capillary surface partitioning, with *H*_*t*_ governing the fraction of the wall area available for direct RBC–tissue exchange (Eq. (21)). At higher hematocrits (*H*_*t*_*≥* 0.5), a large capillary surface fraction is assigned to the direct pathway and the high RBC oxygen content sustains a large driving gradient (*P*_RBC_ *−P*_*T*_ ). Despite the lower MTC, these two combined effects generate a direct flux that more than compensates for the reduction in plasma–tissue exchange, resulting in a marked increase in tissue oxygenation and a corresponding decrease in *P*_RBC_ (Figure 6(a), bottom row). As hematocrit decreases, both effects weaken simultaneously: the direct pathway loses exchange area and driving force. Consequently, below *H*_*t*_ *≈* 0.4, the RBC-tissue pathway can no longer compensate for the reduced plasma–tissue exchange, inverting the trend toward decreased *P*_*T*_ and increased *P*_RBC_. As with the diameter variations, the deviations grow in magnitude along the capillary for all hematocrit values. However, the variations in *P*_*T*_ are considerably larger, reflecting the stronger influence of hematocrit on the capillary surface partitioning.

The contour map (Figure 6(c)) extends these observations to further diameter– hematocrit combinations and reveals that hematocrit is the primary determinant of the sign of Δ*P*_*T*_, while vessel diameter modulates the response as a secondary parameter. The contour lines are denser for small diameters, indicating that Δ*P*_*T*_ is more sensitive to hematocrit in narrow vessels. At low hematocrit values (*H*_*t*_ ≲ 0.25), Δ*P*_*T*_ remains negative across the entire diameter range, with the largest reductions (approximately 60%) occurring in the smallest vessels. The transition between negative and positive deviations in Δ*P*_*T*_ shifts toward lower hematocrit values as the diameter increases, e.g. to *H*_*t*_ *≈* 0.4 for *D* = 4 *µ*m and to *H*_*t*_ *≈* 0.3 for *D ≥* 5 *µ*m, consistent with the parametric results described above. At higher hematocrit values (*H*_*t*_ ≳ 0.5), mean Δ*P*_*T*_ becomes positive, reaching a maximum of approximately +60% for high hematocrits and small diameters (*H*_*t*_ *≈* 0.8*−* 0.9, *D≈* 3 *µ*m), comparable in magnitude to the largest reductions.

Overall, the results suggest that hematocrit primarily determines the sign of the discrepancy between the two formulations, while vessel diameter impacts its magnitude. Small capillaries represent the most sensitive configuration, where the predicted mean *P*_*T*_ can differ by several tens of percent.

In realistic MVNs, the comparison between the single-and double-flux models is performed on a single representative microvascular subnetwork. In contrast to the single-vessel analysis, where the constant MTC (*K*_C_) was used, here the RBC–plasma exchange is described in both models by the Vadapalli formulation, which we select as the reference for the single-flux model. As in the single-vessel case, the differences between the two models therefore reflect only the contribution of the additional RBC– tissue pathway, whose MTC is still based on *D*_RBC_ (Online Resource 1, Table S4). The global distributions only show relatively small differences for the two models (Fig. 7(a); Online Resource 1, Fig. S3). This is consistent with the network characteristics, with a mean diameter of approximately 4 *µ*m and a mean hematocrit of approximately 0.31 (Online Resource 1, Table S1), which place the network near the transition zone of the contour map (Figure 6(c)) where the two formulations yield minimal differences. While the differences between the two distributions are basically negligible for *P*_*pl*_ and *P*_RBC_, the EAT rises from 4.8 to 5.5 mmHg for the double-flux model. Although these differences remain small at the global scale, local pairwise differences in *P*_*T*_ (Fig. 7(b)) are slightly larger, reaching about 3 mmHg.

**Fig. 7.**
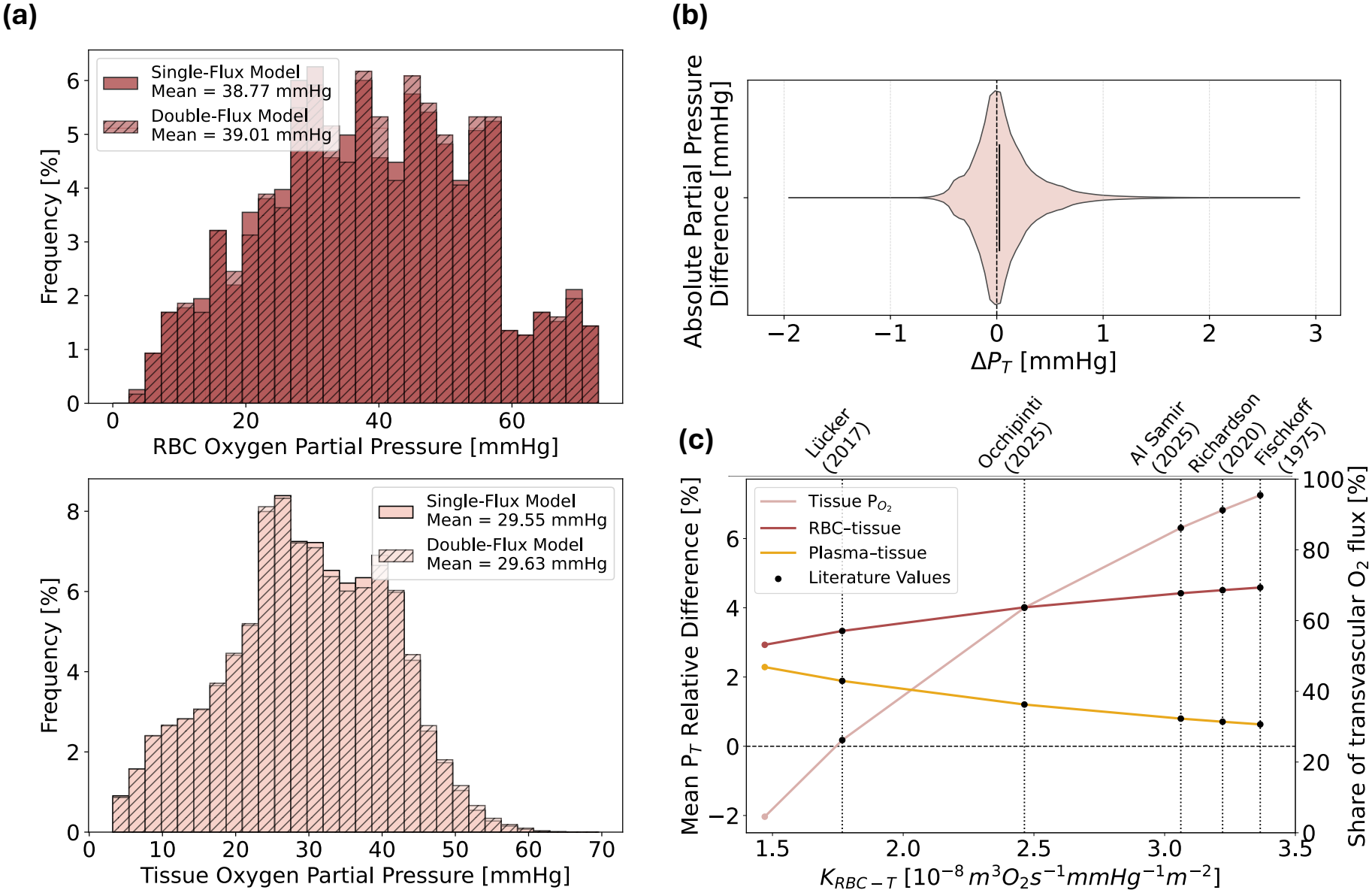
Impact of the double-flux model in realistic microvascular networks and sensitivity to the RBC–tissue mass transfer coefficient *K*_RBC*−*T_. **(a)** Frequency distributions of *P*_*RBC*_ (top) and *P*_*T*_ (bottom) across a representative microvascular subnetwork for the single- and double-flux model. **(b)** Violin plot of pairwise differences in *P*_T_, between the double- and single-flux models. **(c)** Mean relative difference in tissue oxygen partial pressure (*P*_T_, pink, left axis) between the double-and single-flux models, and share of the transvascular oxygen flux (right axis) delivered through the RBC–tissue (dark red) and plasma–tissue (yellow) pathways, as a function of the RBC–tissue mass transfer coefficient *K*_RBC*−*T_. Black markers and dotted vertical lines correspond to *K*_RBC*−*T_ values derived from diffusion coefficients *D*_RBC_ reported in the literature and adjusted to a reference temperature of 37^*°*^C. In all panels, the RBC–plasma mass transfer coefficient is given by the Vadapalli formulation in both the single-and double-flux models. The *K*_RBC*−*T_ value used in realistic MVN simulations (panel (a),(b)) corresponds to the one indicated as Lücker (2017)

#### 3.2.2 Sensitivity of Double-Flux Oxygen Transport to the RBC–Tissue Diffusion Coefficient

With the current set of mass transfer coefficients, the two formulations yield nearly identical global predictions of tissue oxygenation in the realistic network (Fig. 7(a), bottom). However, the robustness of this observation depends on the value assigned to *D*_RBC_, which directly affects the definition of the RBC–tissue MTC according to Eq.(24) and remains highly uncertain. Reported values in the literature span two orders of magnitude, reflecting differences in experimental conditions, temperature, and mod-elling assumptions (see Table 1). Therefore, it is necessary to investigate whether the observed effects persist across the range of diffusion coefficients reported in the literature. To this end, we performed a sensitivity analysis to assess the impact of RBC diffusivity on oxygen transport and on the resulting tissue oxygenation patterns. The analysis was performed on the same representative realistic microvascular network considered in the previous section. The single-flux model with the Vadapalli formulation [37] serves as the reference, against which the double-flux model is evaluated over the range of *D*_RBC_ values reported in the literature, while keeping all other model parameters unchanged.

**Table 1.** Literature-based values of the oxygen diffusion coefficient across the RBC membrane (*D*_RBC_) and corresponding effective RBC–tissue mass transfer coefficients (*K*_RBC−T_). Diffusion coefficients are taken from the cited studies or, where indicated (^*a*^), derived via the Meyer–Overton law, and are adjusted when necessary to a reference temperature of 37^*°*^C using the Arrhenius relation (^*b*^). The resulting *K*_RBC−T_ values are used in the sensitivity analysis of the double-flux model.

| Reference | *D*<sub>RBC</sub><br>[m<sup>2</sup>s<sup>−1</sup>] | **Temperature**<br>[℃] | *D*<sub>RBC</sub> **at** 37℃<br>[m<sup>2</sup>s<sup>−1</sup>] | *K*<sub>RBC−T</sub><br>[m<sup>3</sup> O<sub>2</sub> s<sup>−1</sup> mmHg<sup>−1</sup> m<sup>−2</sup>] |
|---|---|---|---|---|
| Fischkoff [83] | 7.27 · 10<sup>−10</sup> | 37 | 7.27 · 10<sup>−10</sup> | 3.37 · 10<sup>−8</sup> |
| Lücker [35] | <sup>*a*</sup>7.10 · 10<sup>−12</sup> | 37 | 7.10 · 10<sup>−12</sup> | 1.77 · 10<sup>−8</sup> |
| Richardson [82] | 7.00 · 10<sup>−11</sup> | 23 | <sup>*b*</sup>1.21 · 10<sup>−10</sup> | 3.22 · 10<sup>−8</sup> |
| Occhipinti [80] | 5.77 · 10<sup>−12</sup> | 10 | <sup>*b*</sup>1.73 · 10<sup>−11</sup> | 2.46 · 10<sup>−8</sup> |
| Al Samir [81] | 6.00 · 10<sup>−11</sup> | 37 | 6.00 · 10<sup>−11</sup> | 3.06 · 10<sup>−8</sup> |

<sup>*a*</sup> Value derived from *K*<sub>cell</sub>, *α*<sub>pl</sub>, and *σ*<sub>RBC</sub> (Online Resource 1, Table S4) via the Meyer–Overton law [78].
<sup>*b*</sup> Value derived from the Arrhenius relation [84]: $\frac{D_2}{D_1} = \exp\left[\frac{E_a}{R}\left(\frac{1}{T_1} - \frac{1}{T_2}\right)\right]$, where *R* is the universal gas constant (8.314 J mol<sup>−1</sup>K<sup>−1</sup>) and *E*<sub>*a*</sub> is the activation energy for O<sub>2</sub> diffusion in RBC membranes (29.7 kJ mol<sup>−1</sup> [85]).

A monotonic increase in the mean relative difference in *P*_T_ is observed as *K*_RBC*−*T_ increases (Fig. 7(c)). The tissue 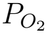 transitions from slightly negative values (approximately *−*2%) at the lower end of the range to approximately +7% at the upper end. It crosses zero near *K*_RBC*−*T_ *≈* 1.75 *×* 10^*−*8^ m^3^ O_2_ s^*−*1^ mmHg^*−*1^ m^*−*2^, which corresponds to the estimate derived from *K*_*C*_ based on values from Lücker et al. [35], in full agreement with the results of the previous section. In contrast, the partitioning of the transvascular oxygen flux between the two pathways changes markedly, with the share delivered directly by the RBCs rising from approximately 53% to 69% over the explored range (about 57% at the zero crossing) at the expense of the plasma–tissue pathway. Above the Lücker estimate, for the higher *K*_RBC*−*T_ values reported by Occhipinti [80], Al Samir [81], Richardson [82], and Fischkoff [83], the double-flux model predicts progressively larger gains in tissue 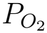, accompanied by a growing contribution of the direct RBC– tissue pathway. Overall, these results underline that a reliable characterization of this parameter is essential for accurate estimates of tissue oxygenation.

### 3.3 Model Validation

Based on our previous investigations and the remaining uncertainties about the diffusion coefficient between RBC–tissue, the Vadapalli model is chosen for validation against in vivo data. The validation study was performed on four realistic microvascular subnetworks, two extracted from MVN1 and two from MVN2. As erythrocyte-associated transients (EATs, Fig. 8(a)) are a hallmark of capillary oxygen transport [18, 38–43] and can be captured by our model through its explicit distinction between RBC and plasma oxygenation, they represent a key quantity of interest for model validation. Here, EATs are computed from the difference between *P*_RBC_ and *P*_pl_. The two variables show a strong correlation, reflecting the tight coupling between the two intravascular compartments (Fig. 8(e)). *P*_RBC_ consistently exceeds *P*_pl_, causing a systematic deviation from the identity line (*P*_RBC_ = *P*_pl_). This reflects the persisting oxygen gradient from RBC to plasma and underlines the model’s ability to capture intracapillary oxygen heterogeneity.

**Fig. 8.**
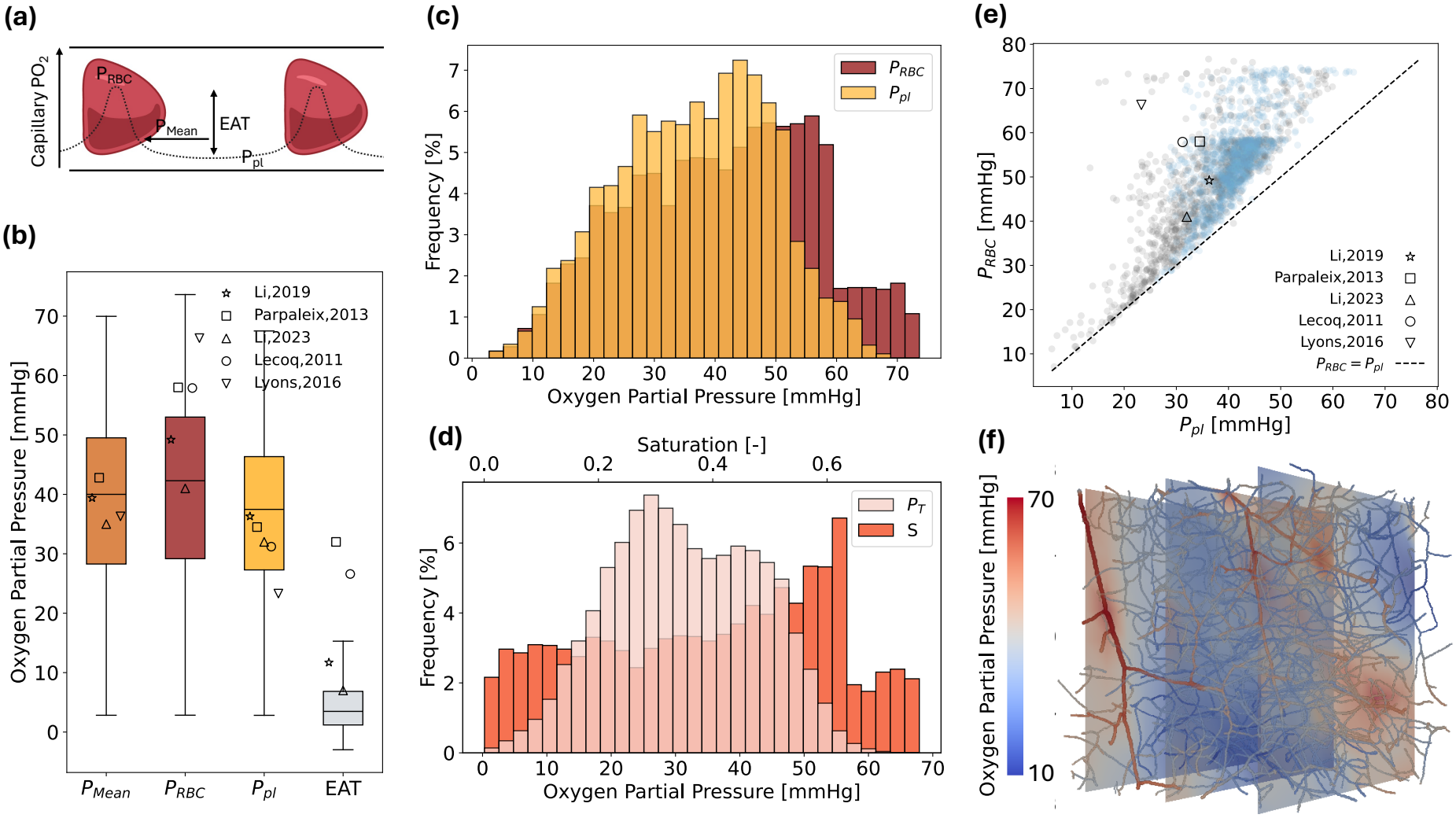
Validation of the oxygen transport model against in vivo experimental data. **(a)** Schematic representation of the intracapillary oxygen distribution. The mean capillary oxygen partial pressure (*P*_*Mean*_) is bounded by the oxygen partial pressure within red blood cells (*P*_*RBC*_) and the lower value in the inter-RBC plasma region (*P*_*pl*_). Their difference defines the erythrocyte-associated transient (EAT).**(b)** Boxplots of the distributions of *P*_*Mean*_, *P*_*RBC*_, *P*_*pl*_, and EAT. Symbols indicate experimental measurements reported in the literature [18, 40–43]. **(c)** Frequency distributions of *P*_*RBC*_ and *P*_*pl*_. **(d)** Frequency distributions of tissue oxygen partial pressure (*P*_*T*_ ) and hemoglobin oxygen saturation (*S*). Results in (b)–(d) are pooled across four realistic microvascular subnetworks. **(e)** Relationship between *P*_*RBC*_ and *P*_*pl*_ for two microvascular networks. The dashed line denotes the identity line (*P*_*RBC*_ = *P*_*pl*_). Symbols indicate experimental measurements reported in the literature. **(f)** Representative microvascular network extracted from MVN1 embedded in 0.064 mm^3^ of cerebral tissue, showcasing oxygen partial pressure distribution, both in vasculature and tissue

The simulated *P*_T_ exhibits a mean value of 31.47 ± 11.49 mmHg, consistent with in vivo imaging measurements [20, 21, 65, 86–88] and previous computational studies [27, 31, 32, 46, 47, 65] (Fig. 8(d)). The distributions predicted for *P*_RBC_, *P*_Mean_ and *P*_pl_ (Fig. 8(b)-(c)) are consistent with experimental observations, reproducing the same qualitative trends observed across independent studies [18, 40–43]. Nevertheless, the resulting mean EAT of 4.53± 4.30 mmHg remains lower than the values typically measured in vivo.

## 4 Discussion

This study introduces a mixed-dimensional 1D–3D computational framework for simulating oxygen transport in anatomically realistic cerebral microvascular networks embedded within parenchymal tissue. The main objective was to characterize oxygen exchange mechanisms at the microvascular scale and to investigate how different descriptions of intravascular oxygen transport influence tissue oxygenation predictions. In contrast to most existing models, which represent blood oxygenation as a single lumped variable [27, 31, 32, 46, 47], our formulation explicitly distinguishes between oxygen dissolved in plasma, oxygen dissolved within RBCs, and oxygen bound to hemoglobin. This phase-resolved description is consistent with experimental observa-tions obtained from in vivo measurements in the mouse brain cortex [18, 38–43]. It also enables the investigation of a double-flux formulation, in which oxygen can reach the tissue through both plasma-mediated transport and direct RBC–tissue exchange, allowing us to assess the validity of the simplified single-flux assumption.

Our study shows that the RBC-plasma MTCs impact local oxygen transport, particularly under conditions of low hematocrit with differences in predicted oxygen partial pressures reaching up to 8 mmHg. At intermediate and high hematocrit levels (*H*_*d*_*∈* [0.2, 1)), the impact of the selected MTC formulations remains small, suggesting that a constant MTC likely provides a reasonable approximation. This behavior reflects the underlying physical mechanism captured by the hematocrit-dependent formulations. In capillaries with single-file flow, hematocrit determines the axial spacing between consecutive RBCs. At high hematocrit, the inter-RBC distance is small and the plasma gaps between neighbouring RBCs are short, so that the oxygen released by the RBCs reaches the capillary wall over short diffusion paths, increasing the efficiency of oxygen transfer. Conversely, at low hematocrit, the larger inter-RBC distance results in longer plasma diffusion pathways and reduced oxygen discharge efficiency. In consequence, hematocrit influences oxygen delivery through two interconnected mechanisms: a reduction in oxygen-carrying capacity due to decreased hemoglobin concentration and a reduction in microvascular oxygen exchange efficiency due to the increased inter-RBC distance. The results further highlight the central role of hematocrit in regulating local oxygen delivery [35], underscoring the importance of accurately resolving hematocrit heterogeneity in realistic microvascular networks, as local deviations may substantially alter local tissue oxygen availability.

Despite the pronounced local differences between MTC formulations, the overall tissue oxygenation distributions remained remarkably robust. This suggests that for applications focused on macroscopic oxygenation patterns, such as measurements with limited spatial resolution, the influence of the specific RBC–plasma MTC formulation may be negligible. However, neglecting the dependence of the MTC on hematocrit and, to a lesser extent, on saturation may then lead to a systematic overestimation of local oxygen availability and affect whether localized regions are classified as normoxic or near-hypoxic. The Vadapalli formulation, which accounts for both dependencies, offers the most physiologically comprehensive description at no additional computational cost and is therefore recommended for applications where spatially resolved oxygenation is of interest.

Compared with the single-flux model, the double-flux formulation reduced tissue oxygenation at low hematocrit (*H*_*d*_ ≲ 0.3), with reductions of up to approximately 60% in single-vessel simulations, and enhanced it at high hematocrit, with increases up to *∼* +60%. The largest differences in both directions were observed in small capil-laries. This behaviour can be explained by the combined effect of two mechanisms. First, introducing a direct RBC–tissue pathway reduces the effective area available for plasma-mediated oxygen exchange. Second, at low hematocrit, the RBC–tissue pathway is not sufficiently strong to compensate for this reduction, because it operates over a small fraction of the capillary wall and the corresponding oxygen transfer remains limited by the RBC membrane mass transfer coefficient. While this at first might appear like an undesirable condition, it implies that under low hematocrit conditions excessive oxygen unloading from RBCs is prevented, thereby preserving oxygen availability in downstream regions. Nonetheless, in regions where hematocrit is generally low the reduced unloading can also become detrimental and contribute to shortages in tissue oxygenation.

The significant impact of the surface available for oxygen discharge to tissue raises a broader question about how this affects oxygen discharge across vessels of different scales. In larger vessels with multi-file flow, oxygen exchange across the entire vessel surface is likely a good approximation. However, the situation is less clear for vessels with single-file flow where only part of the vessel surface is available for direct oxygen exchange from plasma, while the rest of the vessel surface is in direct contact with RBCs. Here, the key question is: is a direct flux from RBC to tissue possible and what relevance does this pathway have in comparison to the sequential RBC-plasma-tissue discharge? Currently, the most suitable MTC for the RBC–tissue discharge suggests that the direct RBC–tissue pathway tends to modestly enhance oxygen unloading to the surrounding tissue in realistic microvascular networks. However, more data is necessary to solidify this observation. This is also relevant for existing single-flux oxygen models, which commonly assume that the whole capillary surface is available for oxygen exchange [28, 32, 46, 65], an approximation that tends to overestimate oxygen discharge and may consequently mask local shortages in oxygen availability.

These findings carry direct implications for ageing and cerebrovascular disorders, where reduced hematocrit [86] and capillary narrowing or rarefaction [6, 89, 90] are common. As shown above, low hematocrit regimes increase the sensitivity of predicted oxygenation to the RBC–plasma MTC formulation, where constant assumptions may lead to an underestimation of hypoxic regions. Similarly, reduced capillary diameters amplify the differences between single-and double-flux models, with potentially significant consequences for oxygen delivery efficiency. A further consequence of simplified model assumptions is the attenuation of tissue oxygenation heterogeneity. Blood flow in the microcirculation is intrinsically heterogeneous [91, 92], and low-flow vessels may approach the hypoxic threshold even under physiological conditions [39, 41]. Both variable RBC–plasma MTC formulations and direct RBC–tissue exchange predict reductions in tissue oxygenation in low-hematocrit, small-diameter regions, suggesting that simplified formulations may underestimate oxygen deficits in the most vulnerable segments of the network. Under pathological conditions that further amplify flow heterogeneity, this limitation is expected to become increasingly severe. These considerations extend beyond the diseased brain: brain regions such as the hippocampus, characterized by lower microvascular density, smaller vessel diameters, and lower hematocrits [93], correspond precisely to the local conditions where the susceptibility to oxygen shortages may be increased and an inappropriate formulation could mask physiologically relevant differences in oxygen availability across brain regions.

The results also inform the interpretation of local regulatory responses to oxygen demand. The single-flux, variable-MTC results indicate that local increases in hematocrit may be particularly effective in low-hematocrit vessels, where the dominant limitation arises from interfacial exchange rather than from convective oxygen supply alone. In this regime, increasing the hematocrit may therefore confer a dual benefit by enhancing convective oxygen transport while simultaneously increasing the MTC. However, beyond a threshold hematocrit (*H*_*d*_ *≈*0.3), further increases result in progressively diminishing gains in tissue oxygenation, as the capillary wall becomes rate-limiting. The double-flux model adds a further dimension to this picture, suggest-ing that in low-hematocrit vessels (*H*_*d*_ ≲ 0.3), both an increase in hematocrit and a vasodilation of small capillaries (*D ≤* 6 *µ*m) may contribute to alleviating oxygen deficits in the tissue. Together, these results suggest that hematocrit regulation and capillary diameter adjustments represent distinct but interacting levers for modulating local oxygen supply, each most effective in specific microvascular regimes.

Currently, our framework describes steady-state oxygen transport. This assumption is justified by the relatively short characteristic times required for oxygen transport to reach a spatially stationary distribution at the microvascular scale, which are expected to occur within a few seconds. Furthermore, cerebral oxygenation is maintained within a relatively narrow physiological range [91, 94], allowing the oxygen distribution to be considered quasi-stationary in the absence of rapid changes in neuronal activity or blood flow. Extending the formulation toward fully time-dependent simulations will be essential for investigating dynamic oxygen delivery and pathological scenarios, e.g. temporary capillary stalling [95].

A key consideration in modeling oxygen transport in the cerebral microcirculation is whether RBC and plasma oxygenation can be assumed to equilibrate instantaneously. This assumption requires that the characteristic time for RBC–plasma oxygen equilibration (*≈* 70-145 ms [4]) is much shorter than the capillary transit time. While this condition may hold when considering transit through the entire capillary bed (*≈* 0.5–2 s [19, 96–98]), it becomes questionable at the scale of individual capillary segments, where transit times of 50–100 ms (estimated for typical capillary lengths of 50–100 *µ*m and RBC velocities of *∼*1 mm/s) are comparable to the equilibration timescale. Moreover, the cerebral microcirculation is characterized by heterogeneous hematocrit distributions and a wide range of capillary transit times, both of which modulate the degree of RBC–plasma equilibration. In vessels with short transit times or low hematocrit, instantaneous equilibration may overestimate oxygen availability in plasma and consequently the flux delivered to tissue. By treating *P*_*pl*_ and *P*_*RBC*_ as distinct variables, the present model captures the additional resistance at the RBC membrane and the hemoglobin-oxygen dissociation kinetics, yielding a more faithful representation of diffusion limitation within the capillary lumen. Greater detail in this regard can be achieved by models such as that of Lücker et al. [52], that resolve individual RBCs as discrete, moving sources, explicitly capturing intra-RBC oxygen diffusion as well as the axial and radial partial pressure heterogeneity arising from the physical spacing between cells. However, these models come at significantly larger computational costs, thus while highly accurate and relevant at the single-capillary scale, such approaches become impractical for the large-network and parametric studies. The present framework therefore introduces an intermediate option that accounts for hemoglobin-oxygen release kinetics and RBC membrane resistance without resolving individual cells, a trade-off especially suitable when network-scale distributions with microscale accuracy are to be predicted.

As current experimental estimates for the RBC membrane oxygen diffusivity span several orders of magnitude, a more rigorous characterization of this coefficient would be crucial for a more accurate quantification of capillary oxygen discharge. Early studies primarily quantified oxygen diffusion within isolated lipid membranes [83], whereas more recent investigations consider the intact RBC membrane and other factors that may influence oxygen transport, including membrane composition, cellular organization, and the contribution of membrane proteins [81, 82]. Other approaches have combined experimental observations with mathematical modeling to quantify how specific membrane components, such as oxygen-permeable channels, contribute to the overall oxygen permeability of RBCs [80]. Therefore, the reported values do not necessarily represent conflicting estimates of the same property, but rather reflect differences in the experimental systems and methodologies used to characterize oxygen transport across the RBC membrane. However, to identify the most suitable MTC for our model, experimental approaches specifically designed to quantify oxygen diffusion across the intact RBC membrane under physiologically relevant conditions need to be developed. Measurements performed directly on erythrocytes [82, 99], using standardized methodologies and conditions representative of the microvascular environment, could provide more consistent estimates of membrane oxygen diffusivity. Such efforts would reduce the current uncertainty in RBC membrane oxygen transport properties and provide more reliable parameters for computational models of oxygen transport.

In summary, the main limitations of the presented framework are the following. At the level of the model formulation, oxygen transport is described at steady state, therefore transient phenomena such as neurovascular coupling, flow fluctuations, and capillary stalling are not captured. In addition, RBCs are represented as a continuous phase rather than as discrete moving cells, which limits the resolution of intracapillary heterogeneity and may contribute to the underestimation of EATs. A second group of limitations concerns the model parameters: (i) the RBC–plasma MTCs rely on extrapolation of the hematocrit–saturation space, (ii) the RBC–tissue MTC remains highly uncertain, and (iii) all transfer coefficients are assumed to be independent of the local flow. Likewise, the tissue is modeled as a homogeneous medium with spatially uniform oxygen consumption, i.e. neglecting depth-or cell-type-dependent metabolic demand. Finally, some limitations arise from the simulated configurations. The network analysis is limited to four cortical subnetworks of 0.064 mm^3^, whose oxygenation is also influenced by the imposed inflow boundary conditions. Moreover, the zero-flux (homogeneous Neumann) condition imposed on the outer tissue boundary neglects oxygen exchange with the surrounding tissue and may bias tissue PO_2_ near the domain faces. However, while these limitations might slightly impact the resulting oxygen fields, they are not expected to influence our observations related to the different MTC formulations.

Overall, our results demonstrate that accurate representation of intercompartmental mass transfer is essential for predicting physiological oxygen fields across scales. Hematocrit-dependent RBC–plasma transfer formulations prove to be particularly relevant for the heterogeneous flow conditions typical for microvascular flow. Particularly for the smallest vessels, incorporating a direct RBC–tissue flux may improve the physiological representation of oxygen transport, although accurate mass transfer coefficients will be essential to fully realize the potential gains in model accuracy. Taken together, our model provides a versatile computational platform particularly well suited for resolving capillary oxygen discharge and establishes an important foundation for future investigations of cerebral energy homeostasis in both health and disease.

## Supporting information

Online Resource 1

## Acknowledgments

The authors thank Chryso Lambride and Sofia Farina for valuable discussions and feedback on the figures, and Artemisia Sarteschi for her contribution to the implementation of the microBlooM blood flow iterative solver.

## Statements and Declarations

### Supplementary Information

Online Resource 1 (ESM_1.pdf) contains the convergence criteria of the blood flow model, the characteristics of the microvascular sub-networks, the derivation of the intravascular oxygen balance, the boundary conditions, the numerical implementation, the grid convergence study, the model parameters and additional results (Sections S1–S8), together with Supplementary Tables S1–S5 and Supplementary Figs. S1–S3.

### Ethics Approval

Not applicable.

### Data Availability

All data reported in this paper will be shared by the corresponding author upon reasonable request.

### Code Availability

The oxygen transport model developed in this study is publicly available at https://github.com/GaiaStievano/Oxygen-Transport-Model. Blood flow simulations were performed using the open-source microBlooM framework, available at https://github.com/Franculino/microBlooM.

### Author Contributions

Conceptualization: G.S., F.S.; Methodology: G.S., G.B., D.C.P., T.K.; Software: G.S., T.K.; Data curation: G.S.; Formal analysis: G.S., G.B., D.C.P., F.S.; Investigation: G.S., G.B., D.C.P., F.S.; Validation: G.S.; Visualization: G.S., D.C.P.; Writing – original draft: G.S., F.S.; Writing – review & editing: F.S., T.K., G.B.,D.C.P.; Supervision: F.S.; Funding acquisition: F.S. All authors read and approved the final manuscript.

