## Supplementary material for "Mass Transfer Formulations Shape Oxygen Transport and Tissue Oxygenation in Microvascular Networks": Online Resource 1

*Annals of Biomedical Engineering*

Gaia Stievano*[1], Dominic Campbell-Pitt[1], Giorgia Barra[1,2], Timo Koch[3] and Franca Schmid[1]

[1] *Cardiovascular Engineering, ARTORG Center for Biomedical Engineering, University of Bern, Freiburgstrasse 3, 3012 Bern, Switzerland*
[2] *Paris Brain Institute (ICM), Institut du Cerveau, 47 Boulevard de l'Hôpital, 75013 Paris, France*
[3] *Department of Hydromechanics and Modelling of Hydrosystems, University of Stuttgart, Pfaffenwaldring 61, 70569 Stuttgart, Germany*

### S1 Convergence of Iterative Model

#### S1.1 Convergence Criteria

Unlike existing microvascular blood flow models, the convergence of the iterative model used in this study is assessed through a residual-based criterion that enforces RBC mass conservation at the internal nodes of the network. Let $R_j^{(i)}$ denote the residual of the RBC mass balance at node $j \in \mathcal{N}_{\text{int}}$ (set of internal network nodes) at iteration $i$, defined as

$$R_j^{(i)} = \sum_{k \in \mathcal{E}_j} s_{jk} Q_k^{(i)} H_k^{(i)} \tag{S1}$$

where $\mathcal{E}_j$ denotes the set of edges connected to node $j$; $Q_k^{(i)}$ and $H_k^{(i)}$ represent the blood flow rate and hematocrit in a given edge $k \in \mathcal{E}_j$, respectively, and $s_{jk} = \pm 1$ indicates the direction of flow (inflow/outflow) relative to the node $j$. The global residual at iteration $i$ is defined as the maximum nodal residual:

$$R_{\max}^{(i)} = \max_{j \in \mathcal{N}_{\text{int}}} \left| R_j^{(i)} \right| \tag{S2}$$

Convergence is achieved when this residual falls below a prescribed threshold $\varepsilon$:

$$R_{\max}^{(i)} \leq \varepsilon \tag{S3}$$

To account for the wide range of flow magnitudes in microvascular networks, the convergence threshold $\varepsilon$ is scaled according to the characteristic magnitude of the specific flow problem under investigation. Specifically,

$$\varepsilon = 10^{3+M}\ \text{m}^3\ \text{s}^{-1}, \tag{S4}$$

where $10^M$ is the order of magnitude of the maximum blood flow residual at the first iteration ($i = 0$):

$$R_{Q,\max}^{(0)} = \max_{j \in \mathcal{N}_{\text{int}}} \left| \sum_{k \in \mathcal{E}_j} s_{jk} Q_k^{(0)} \right| \tag{S5}$$

The problem-dependent scaling of the convergence threshold ensures numerical stability and strict RBC mass conservation across networks with different flow magnitudes. Additionally, a limit of $i_{\max} = 4000$ iterations is imposed to ensure computational efficiency; exceeding this limit classifies the solution as non-convergent. This condition was not encountered in the present study, as all simulated networks achieved convergence within the prescribed tolerance.

#### S1.2 Successive Over-Relaxation Scheme

Approaches to improve convergence reported in the literature [1, 2] commonly rely on the successive over-relaxation (SOR) method to accelerate convergence and improve numerical stability. The SOR scheme is applied directly to the hematocrit update, which represents the primary variable of interest in the iterative procedure. Let $H^{(i)}$ denote the hematocrit at iteration $i$ and $\tilde{H}^{(i+1)}$ the predicted value at iteration $i+1$. The updated solution $H^{(i+1)}$ is computed as a weighted average:

$$H^{(i+1)} = (1 - \alpha^{(i+1)}) H^{(i)} + \alpha^{(i+1)} \tilde{H}^{(i+1)}, \tag{S6}$$

where $\alpha \in [0, 1]$ is the relaxation factor. The relaxation factor may be kept constant or updated during the iterations. In the present implementation, $\alpha$ is initially set to 1 and reduced by 10% every 50 iterations. For the two microvascular networks considered in this study (MVN1 and MVN2), convergence is reached in 705 and 685 iterations, with final relaxation factors of $\alpha = 0.23$ and $\alpha = 0.25$.

### S2 Microvascular Subnetworks

Four subnetworks, two extracted from MVN1 and two from MVN2, were used for the oxygen transport simulations in realistic microvascular networks (Section 2.2 of the main text). Their structural and hemodynamic characteristics are summarized in Table S1.

| Subnetwork ID | MVN | No. DA | No. AV | No. C | Mean Hematocrit | Mean Blood Flow $[\mathrm{m}^3/\mathrm{s}]$ | Mean Diameter $[\mu\mathrm{m}]$ | Total Vessel Length [mm] |
|---|---|---|---|---|---|---|---|---|
| 1 | 1 | 17 (2) | 0 (0) | 1183 | 0.32 | $9.48 \times 10^{-15}$ | 3.92 | 62.81 |
| 2 | 1 | 11 (2) | 15 (2) | 1073 | 0.31 | $8.30 \times 10^{-15}$ | 3.91 | 57.49 |
| 3 | 2 | 15 (1) | 40 (2) | 1229 | 0.32 | $7.57 \times 10^{-15}$ | 4.26 | 63.60 |
| 4 | 2 | 20 (2) | 26 (1) | 1347 | 0.31 | $1.77 \times 10^{-14}$ | 4.07 | 69.78 |

DA: Descending Arterioles; AV: Ascending Venules; C: Capillaries. For DA and AV, values indicate the number of segments within each subnetwork, with the number in parentheses indicating how many distinct parent trees in the original MVN they belong to. Each subnetwork is embedded in a tissue volume of $0.064\ \mathrm{mm}^3$.

**Table S1** Structural and hemodynamic characteristics of the analyzed realistic microvascular subnetworks.

### S3 Intravascular Oxygen Balance

The total oxygen flux through a vessel section is the sum of three contributions: oxygen dissolved in plasma, oxygen dissolved in RBCs, and oxygen bound to hemoglobin. Since $(1 - H_d)A_v u$ and $H_d A_v u$ are the plasma and RBC fluxes, and a unit volume of RBCs carries an amount $C_0 S$ of bound oxygen, the balance of the total intravascular oxygen per unit vessel length reads

$$\frac{\partial}{\partial s}\Big( A_v u \left[(1 - H_d)\, \alpha_{pl} P_{pl} + H_d\, \alpha_{RBC} P_{RBC} + H_d\, C_0 S\right] \Big) = \hat{q}_C, \tag{S7}$$

i.e. the total oxygen flux changes along a vessel only through the exchange with the tissue. Writing a separate balance for each of the three contributions yields Eqs. (2) and (3) of the main text for the oxygen dissolved in plasma and in RBCs, and, for the oxygen bound to hemoglobin,

$$\frac{\partial}{\partial s}\left(C_0 H_d A_v u\, S\right) + C_0 H_d A_v F(P_{RBC}, S) = 0. \tag{S8}$$

Since $C_0$ is constant, it can be simplified in Eq. (S8). Moreover, along each vessel segment both the blood flow rate $A_v u$ and the RBC flux $H_d A_v u$ are conserved,

$$\frac{\partial}{\partial s}(A_v u) = 0, \qquad \frac{\partial}{\partial s}(H_d A_v u) = 0, \tag{S9}$$

so that $H_d$ can be taken out of the derivative and simplified (for $H_d > 0$), which yields Eq. (4) of the main text.

At bifurcations, $H_d$ changes as a result of phase separation, and the conservation of bound oxygen must be imposed explicitly. Let $\mathcal{I}(n)$ and $\mathcal{O}(n)$ denote the segments flowing into and out of node $n$, $Q_k = A_{v,k} u_k$ and $S_k$ the flow rate and saturation of segment $k$ at the node, and $S_n$ the saturation in the edge leaving the node. Conservation of the bound-oxygen flux gives $\sum_{k\in\mathcal{I}(n)} H_{d,k} Q_k S_k = S_n \sum_{j\in\mathcal{O}(n)} H_{d,j} Q_j$, and since RBCs are conserved at the node, $\sum_{j\in\mathcal{O}(n)} H_{d,j} Q_j = \sum_{k\in\mathcal{I}(n)} H_{d,k} Q_k$, so that

$$S_n = \frac{\sum_{k\in\mathcal{I}(n)} Q_k H_{d,k}\, S_k}{\sum_{k\in\mathcal{I}(n)} Q_k H_{d,k}}, \tag{S10}$$

which every segment in $\mathcal{O}(n)$ takes as its inflow value. If all inflow segments are RBC-free ($H_d = 0$), the denominator of Eq. (S10) vanishes and the weights $Q_k$ are used instead. The same balance is applied to the dissolved oxygen, with weights $Q_k H_{d,k}$ for $P_{RBC}$ and $Q_k(1 - H_{d,k})$ for $P_{pl}$, so that the total oxygen flux of Eq. (S7) is conserved at every node.

### S4 Boundary Conditions

Inflow values in the realistic MVN are not uniform but depend on vessel type, namely descending arterioles (DA), ascending venules (AV), and capillaries (C). In particular, the inflow RBC oxygen partial pressure in capillaries is defined as a function of the discharge hematocrit $H_d$. This choice accounts for the heterogeneous distribution of RBCs in the microcirculation and the resulting variability in oxygen delivery. Specifically, the capillary inflow value $P_{RBC}^{\text{inflow}}$ is defined through a piecewise function $f(H_d)$, based on experimental data reported by Lyons et al. [3]:

$$f(H_d) = \begin{cases} 48 & H_d \le 0.2 \\ 59 & 0.2 < H_d \le 0.4 \\ 66 & 0.4 < H_d \le 0.6 \\ 70 & H_d > 0.6 \end{cases} \tag{S11}$$

This formulation allows the model to capture the dependence of oxygen availability on hematocrit. The parameter values used to compute the inflow fluxes are summarized in Table S2.

### S5 Numerical Implementation

The monolithic nonlinear system introduced in Section 2.4 of the main text is solved using Newton's method. At each Newton iteration $i$, the following linear system is solved:

$$\mathbf{J}(\mathbf{u}^i)\,\Delta\mathbf{u}^i = -\mathbf{r}(\mathbf{u}^i), \tag{S12}$$

with

$$\mathbf{u} = \begin{bmatrix} \mathbf{u}_v \\ \mathbf{u}_t \end{bmatrix}, \quad \mathbf{J} = \begin{bmatrix} \mathbf{J}_{vv} & \mathbf{J}_{vt} \\ \mathbf{J}_{tv} & \mathbf{J}_{tt} \end{bmatrix}, \quad \Delta\mathbf{u} = \begin{bmatrix} \Delta\mathbf{u}_v \\ \Delta\mathbf{u}_t \end{bmatrix}, \quad \mathbf{r} = \begin{bmatrix} \mathbf{r}_v \\ \mathbf{r}_t \end{bmatrix}. \tag{S13}$$

Here, $\mathbf{r}(\mathbf{u})$ denotes the nonlinear residual function, which depends on the vector of primary unknowns $\mathbf{u}$. The subscripts $v$ and $t$ respectively denote the vascular and

| Parameter | Description | Domain | Value | Reference | Units |
|---|---|---|---|---|---|
| $P_{pl}^{\text{inflow}}$ | | | | | |
| | Inflow Plasma $PO_2$ | $\partial\Lambda_I$ | 25 | [3, 4] | mmHg |
| $P_{RBC}^{\text{inflow}}$ | | | | | |
| | Inflow RBC $PO_2$ | DA $\in \partial\Lambda_I$ | 75 | [3–7] | mmHg |
| | Inflow RBC $PO_2$ | AV $\in \partial\Lambda_I$ | 40 | [3–7] | mmHg |
| | Inflow RBC $PO_2$ | C $\in \partial\Lambda_I$ | $f(H_d)$ | [a] | mmHg |
| $S^{\text{inflow}}$ | | | | | |
| | Inflow S | DA $\in \partial\Lambda_I$ | 0.76 | [b] | - |
| | Inflow S | AV $\in \partial\Lambda_I$ | 0.38 | [b] | - |
| | Inflow S | C $\in \partial\Lambda_I$ | $S(f(H_d))$ | [b] | - |

DA: Descending Arterioles; AV: Ascending Venules; C: Capillaries.

[a] The function $f(H_d)$ is defined in Eq. (S11), based on data extracted from the work of Lyons et al. [3]

[b] Saturation inflow values are set in equilibrium with $P_{RBC}^{\text{inflow}}$ via Eq. (7) of the main text

**Table S2** Inflow boundary conditions prescribed in the network simulations. RBC oxygen partial pressure and hemoglobin saturation are imposed according to vessel type, with capillary inlet values determined as functions of discharge hematocrit.

tissue domains. $\mathbf{J}$ is the Jacobian matrix, which can be split into four different blocks: $\mathbf{J}_{vv} = \partial\mathbf{r}_v/\partial\mathbf{u}_v$ associated with the vascular equations, $\mathbf{J}_{tt} = \partial\mathbf{r}_t/\partial\mathbf{u}_t$ associated with the tissue equations, $\mathbf{J}_{vt} = \partial\mathbf{r}_v/\partial\mathbf{u}_t$ representing the coupling from tissue to the vascular domain, and $\mathbf{J}_{tv} = \partial\mathbf{r}_t/\partial\mathbf{u}_v$ representing the coupling from the vascular to the tissue domain. $\mathbf{\Delta u}$ denotes the Newton correction, i.e. the difference of the current solution to the solution of the previous iteration. To ensure physical admissibility of the solution at each iteration, the Newton update is followed by a component-wise projection $\mathcal{P}$ onto the set of admissible values:

$$\mathbf{u}^{i+1} = \mathcal{P}\left(\mathbf{u}^i + \Delta\mathbf{u}^i\right), \tag{S14}$$

where $\mathcal{P}$ is defined component-wise as:

$$\mathcal{P}(u) = \begin{cases} \min\left(\max(u,\, 0),\, 1\right) & \text{if } u = S, \\ \min\left(\max(u,\, 0),\, 200\right) & \text{if } u = P_{\text{RBC}}, \\ \max(u,\, 0) & \text{if } u = P_k,\ k \in \{\text{pl}, T\}. \end{cases} \tag{S15}$$

This enforces $S \in [0, 1]$, non-negative partial pressures in all compartments, and the upper bound of $200$ mmHg on $P_{\text{RBC}}$. The iterations are repeated until convergence is achieved, with the maximum relative shift between successive Newton iterates required to fall below $10^{-7}$. The linear systems arising at each Newton iteration are solved using a left-preconditioned stabilized biconjugate gradient method (BiCGStab), with a block-diagonal ILU0 preconditioner [8] [9].

### S6 Numerical Convergence Study

Since the present model is based on a coupled 1D–3D formulation, the numerical convergence of the vascular and tissue domains must be considered separately. The vascular network is described through a graph representation in which each vessel (edge) is defined by a sequence of intermediate points connecting its terminal nodes. These points preserve the vessel tortuosity and geometric characteristics, resulting in segment lengths that are not necessarily uniform. In the present implementation, the vascular resolution, with segment lengths ranging from $0.5$ to $1.7\mu$m, is finer than the spatial resolution of the tissue domain. Therefore, the vascular discretization is not expected to be the limiting factor for the numerical accuracy of the coupled model. The grid convergence study consequently focuses on the 3D tissue domain, with the aim of identifying an optimal Cartesian grid resolution that provides an appropriate balance between computational cost and accuracy in predicting tissue oxygen partial pressure. Since no analytical solution is available for this problem, a reference solution computed on a very fine spatial resolution is used for the grid convergence study. The errors of coarser grids are then evaluated by mapping their solutions onto this reference grid.

#### S6.1 Grid Hierarchy

We consider a sequence of computational grids with different spatial resolutions. Let $\mathcal{M}_h$ denote a structured Cartesian grid with characteristic mesh size $h$. The grid $\mathcal{M}_h$ is defined as the union of all control volumes (grid cells) $K$ belonging to that grid $\mathcal{M}_h = \bigcup_{K\in\mathcal{M}_h} K$, and the number of elements of the grid is denoted by $|\mathcal{M}_h|$. Similarly, the reference grid with the finest resolution $h_{\text{ref}}$ is denoted by $\mathcal{M}_{h_{\text{ref}}}$ and is defined as $\mathcal{M}_{h_{\text{ref}}} = \bigcup_{L\in\mathcal{M}_{h_{\text{ref}}}} L$, where $|\mathcal{M}_{h_{\text{ref}}}|$ is the number of elements of the reference grid. The sequence of grids is constructed by successive refinement such that the mesh size doubles between two consecutive grid levels: $h = 2^n h_{\text{ref}}$, with $n \in \{0, 1, 2, 3, 4, 5\}$. With this refinement strategy, each element of a coarser grid contains exactly eight elements of the next finer grid. Therefore, for every element $L \in \mathcal{M}_{h_{\text{ref}}}$, there exists exactly one element $K \in \mathcal{M}_h$ such that $L \subset K$. This hierarchical structure allows for a consistent comparison between solutions computed on different grids.

#### S6.2 Mapping Between Grids

Let $u_h \in \mathbb{R}^{|\mathcal{M}_h|}$ denote the numerical solution on the coarse grid $\mathcal{M}_h$, and let $u_{\text{ref}} \in \mathbb{R}^{|\mathcal{M}_{h_{\text{ref}}}|}$ be the reference solution computed on the finest grid $\mathcal{M}_{h_{\text{ref}}}$, with $h_{\text{ref}} < h$. To enable a consistent comparison between solutions defined on different grids, all quantities are evaluated on the reference grid. To this end, we introduce the interpolation operator

$$\mathcal{I}_{h\to h_{\text{ref}}} : \mathbb{R}^{|\mathcal{M}_h|} \to \mathbb{R}^{|\mathcal{M}_{h_{\text{ref}}}|},$$

which maps the coarse-grid solution onto the fine grid. The interpolated coarse-grid solution evaluated at a fine-grid cell center $\mathbf{x}_f$ is defined as

$$\tilde{u}_h(\mathbf{x}_f) := \left(\mathcal{I}_{h\to h_{\text{ref}}} u_h\right)(\mathbf{x}_f), \quad \forall \mathbf{x}_f \text{ cell center of } L \in \mathcal{M}_{h_{\text{ref}}}.$$

Since $\mathcal{M}_h$ is a structured Cartesian grid, each point $\mathbf{x}_f = (x_f, y_f, z_f)$ lies within a unique coarse-grid cell. Therefore, there exist indices $i, j, k$ such that

$$x_i \le x_f \le x_{i+1}, \quad y_j \le y_f \le y_{j+1}, \quad z_k \le z_f \le z_{k+1},$$

where $(x_i, y_j, z_k) \in \mathcal{M}_h$ denote neighboring coarse-grid cell centers. The interpolated value $\tilde{u}_h(\mathbf{x}_f)$ is obtained via trilinear interpolation:

$$\tilde{u}_h(\mathbf{x}_f) = \sum_{\alpha=0}^{1}\sum_{\beta=0}^{1}\sum_{\gamma=0}^{1} w_{\alpha\beta\gamma}(\mathbf{x}_f)\, u_{h,\, i+\alpha,\, j+\beta,\, k+\gamma},$$

with weights

$$w_{\alpha\beta\gamma}(\mathbf{x}_f) = (1-\xi)^{1-\alpha}\xi^{\alpha}\,(1-\eta)^{1-\beta}\eta^{\beta}\,(1-\zeta)^{1-\gamma}\zeta^{\gamma},$$

and local coordinates

$$\xi = \frac{x_f - x_i}{x_{i+1} - x_i}, \quad \eta = \frac{y_f - y_j}{y_{j+1} - y_j}, \quad \zeta = \frac{z_f - z_k}{z_{k+1} - z_k}.$$

The accuracy of the numerical solution $u_h$ is then assessed by comparing the interpolated coarse-grid solution $\tilde{u}_h$ with the reference solution $u_{\text{ref}}$ at each fine-grid cell center. The pointwise error is defined as:

$$e_h(\mathbf{x}_f) = \tilde{u}_h(\mathbf{x}_f) - u_{\text{ref}}(\mathbf{x}_f), \quad \forall \mathbf{x}_f \text{ cell center of } L \in \mathcal{M}_{h_{\text{ref}}}.$$

Global error metrics are computed over all fine-grid cell centers. In particular, the discrete $L^2$ norm is approximated via numerical integration over the reference grid:

$$\|e_h\|_2 \approx \sqrt{\sum_{\mathbf{x}_f \in \mathcal{M}_{h_{\text{ref}}}} \left(e_h(\mathbf{x}_f)\right)^2 \Delta V_{\text{ref}}}, \qquad \|u_{\text{ref}}\|_2 \approx \sqrt{\sum_{\mathbf{x}_f \in \mathcal{M}_{h_{\text{ref}}}} \left(u_{\text{ref}}(\mathbf{x}_f)\right)^2 \Delta V_{\text{ref}}},$$

where $\Delta V_{\text{ref}} = \Delta x_{\text{ref}}\, \Delta y_{\text{ref}}\, \Delta z_{\text{ref}}$ is the volume of a cell in the reference grid. The relative $L^2$ error is then defined as

$$\epsilon_{L^2} = \frac{\|e_h\|_2}{\|u_{\text{ref}}\|_2}.$$

This metric quantifies the discrepancy between the interpolated coarse-grid solution and the reference solution evaluated on the finest grid, with spatial resolution $h_{\text{ref}} = 1\,\mu$m.

The grid convergence analysis confirms that the simulated solution converges towards the reference solution as the mesh is refined (Fig. S1). The oxygen partial pressure in the tissue exhibits an approximately first-order convergence behavior in the $L^2$ norm, with an observed convergence rate of 1.18. This rate is consistent with the expected performance of the adopted TPFA discretization for coupled transport problems with embedded sources and nonlinear exchange terms [8]. The relative $L^2$ error decreases rapidly with mesh refinement and falls below $2\%$ for grid resolutions finer than $8\,\mu\text{m}$ (Table S3). Based on the smooth convergence trend, an intermediate spatial resolution of $h = 2.5\,\mu\text{m}$ is selected for subsequent analyses, as it is expected to provide an accuracy comparable to the $h = 2\,\mu\text{m}$ grid while reducing computational cost.

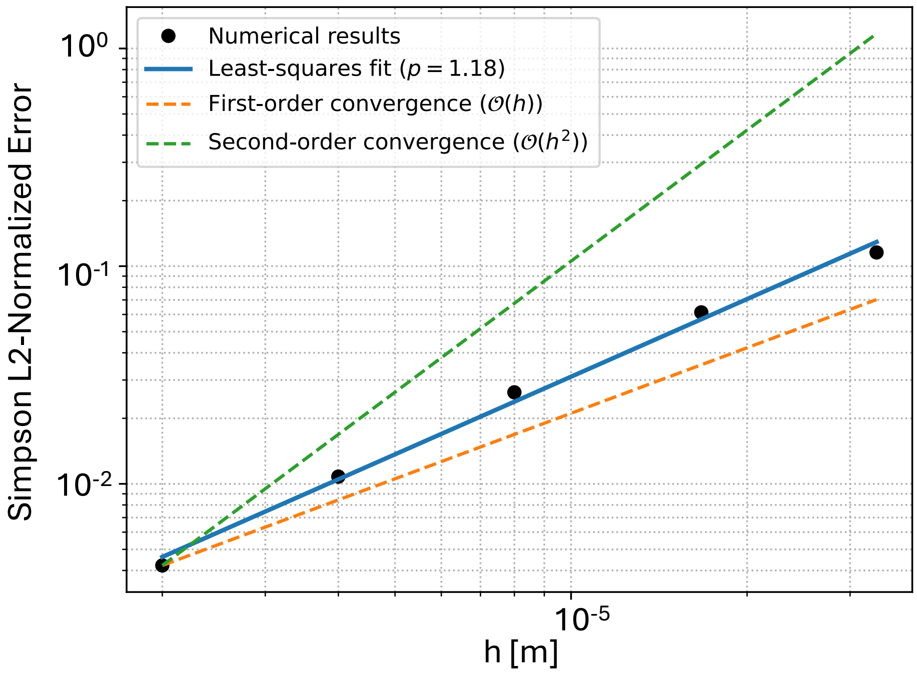


**Fig. S1 Assessment of spatial discretization errors in the 3D tissue domain** Normalized relative $L^2$ error of the tissue oxygen partial pressure solution as a function of the 3D grid cell size $h$. The error is computed with respect to the reference solution obtained on the finest grid ($h_{\text{ref}} = 1\ \mu$m) after interpolation of the coarser-grid solutions onto the reference mesh. Black markers represent the numerical results reported in Table S3. The blue curve corresponds to a least-squares fit of the computed errors, while the orange and green lines indicate the theoretical first-order and second-order convergence rates, respectively.

| 3D Grid Resolution | $h$ [$\mu$m] | DuMuX Time [min] | Relative $L^2$ Error |
|---|---|---|---|
| $200 \times 200 \times 200$ | 1 | 22.8 | - |
| $100 \times 100 \times 100$ | 2 | 3.3 | 0.0042 |
| $50 \times 50 \times 50$ | 4 | 0.42 | 0.0108 |
| $25 \times 25 \times 25$ | 8 | 0.10 | 0.0263 |
| $12 \times 12 \times 12$ | 16.7 | 0.068 | 0.0613 |
| $6 \times 6 \times 6$ | 33.3 | 0.096 | 0.1156 |

**Table S3** Grid convergence analysis for the three-dimensional tissue domain in the coupled 1D–3D framework. The 3D mesh is refined, while the 1D vascular domain is kept fixed with a maximum cell size of $1.7\ \mu$m. The finest grid ($h_{\text{ref}} = 1\ \mu$m) is used as reference for the computation of the relative $L^2$ error. The computational cost is reported in terms of DuMuX runtime. All simulations were performed on a MacBook Pro 14-inch (2021) equipped with an Apple M1 Pro 8-core CPU and 32 GB of RAM.

### S7 Model Parameters

Table S4 lists all the parameters used in the oxygen transport model introduced in Section 2 of the main text, including their definitions, values, units, and sources. When not directly available from the literature, parameters are either estimated or derived as specified in the table.

| Parameter | Description | Value | Units | Reference |
|---|---|---|---|---|
| $\alpha_{pl}$ | $O_2$ solubility in plasma | $2.82 \cdot 10^{-5}$ | $m^3$ $O_2$ $mmHg^{-1}$ $m^{-3}$ | [10] |
| $\alpha_{RBC}$ | $O_2$ solubility in RBCs | $3.38 \cdot 10^{-5}$ | $m^3$ $O_2$ $mmHg^{-1}$ $m^{-3}$ | [11] |
| $\alpha_T$ | $O_2$ solubility in the tissue | $3.89 \cdot 10^{-5}$ | $m^3$ $O_2$ $mmHg^{-1}$ $m^{-3}$ | [12] |
| $\alpha_w$ | $O_2$ solubility in the capillary wall | $3.89 \cdot 10^{-5}$ | $m^3$ $O_2$ $mmHg^{-1}$ $m^{-3}$ | [See $\alpha_T$] |
| $D_T$ | $O_2$ diffusivity in the tissue | $2.41 \cdot 10^{-9}$ | $m^2s^{-1}$ | [13] |
| $D_w$ | $O_2$ diffusivity in the capillary wall | $8.73 \cdot 10^{-10}$ | $m^2s^{-1}$ | [14] |
| $D_{RBC}$ | $O_2$ diffusivity across the RBC membrane | $7.10 \cdot 10^{-12}$ | $m^2s^{-1}$ | [a] |
| $\sigma_w$ | Capillary wall thickness | $1 \cdot 10^{-6}$ | m | [15] |
| $\sigma_{RBC}$ | RBC membrane thickness | $7.5 \cdot 10^{-9}$ | m | [16, 17] |
| $V_{RBC}$ | RBC volume | $4.55 \cdot 10^{-17}$ | $m^3$ | [18] |
| $N_{Hb}$ | Total heme density | 20.3 | mol $m^{-3}$ | [19] |
| $V_{mol,O_2}$ | $O_2$ molar volume at 36.9°C | $2.54 \cdot 10^{-2}$ | $m^3$ $O_2$ $mol^{-1}$ | [20] |
| $C_0$ | Oxygen binding capacity | 0.52 | $m^3$ $O_2$ $m^{-3}$ | [b] |
| $K$ | Dissociation rate constant | 44 | $s^{-1}$ | [19] |
| $n$ | Hill exponent | 2.65 | - | [19] |
| $P_{RBC,50}$ | RBC partial pressure at hemoglobin half saturation | 47.9 | mmHg | [20] |
| $M_0$ | Oxygen consumption | $7 \cdot 10^{-4}$ | $m^3$ $O_2$ $m^{-3}$ $s^{-1}$ | [20, 21] |
| $P_{T,crit}$ | Critical $P_T$ | 1.0 | mmHg | [22] |
| $K_{\text{cell}}$ | RBC–Plasma MTC | $2.67 \cdot 10^{-8}$ | $m^3$ $O_2$ $mmHg^{-1}$ $m^{-2}$ $s^{-1}$ | [c] |
| $H_D^{\text{boundary}}$ | Inflow hematocrit for blood flow initialization | 0.35 | - | [Prescribed] |
| $\mu_b$ | Plasma viscosity | 0.0012 | Pa s | [23] |

[a] Value derived from $K_{\text{cell}}$, $\alpha_{\text{pl}}$, and $\sigma_{\text{RBC}}$ via the Meyer–Overton law [24]

[b] Value derived as: $C_0 = N_{Hb} V_{mol,O_2}$ [20]

[c] Value derived from Eq. (17) of the main text at $H_t = 0.31$ ($K_L(H_t = 0.31)$)

**Table S4** Parameters employed in the oxygen transport model. For each parameter, the adopted value, units, and source are reported. Parameters not directly available from the literature were derived or estimated as indicated.

| Parameter | Description | Value | Units | Reference |
|---|---|---|---|---|
| $P_{pl}^{\text{in}}$ | Inflow plasma $PO_2$ | 25 | mmHg | [3, 4] |
| $P_{RBC}^{\text{in}}$ | Inflow RBC $PO_2$ | 59 | mmHg | [a] |
| $S^{\text{in}}$ | Inflow hemoglobin saturation | 0.63 | – | [b] |
| $J_{\text{in}}/(M_0 V_T)$ | $O_2$ inflow relative to maximal tissue demand | 1.12 | – | [c] |

[a] Value extracted from Lyons et al. [3], corresponding to $0.2 < H_d \leq 0.4$.

[b] Saturation inflow values are set in equilibrium with $P_{RBC}^{\text{in}}$ via Eq. (7) of the main text.

[c] $J_{\text{in}} = Q\left[(1 - H_d)\,\alpha_{pl} P_{pl}^{\text{in}} + H_d\left(\alpha_{RBC} P_{RBC}^{\text{in}} + C_0 S^{\text{in}}\right)\right]$ is the total $O_2$ inflow at the vessel inlet, $V_T$ the volume of the tissue domain, and $M_0$ the maximal $O_2$ consumption rate (Table S4). The ratio is identical for all hematocrit levels.

**Table S5** Inflow boundary conditions prescribed for the single-vessel benchmark and resulting ratio between oxygen inflow and maximal tissue demand. Plasma and RBC oxygen partial pressures are imposed at the vessel inlet, while hemoglobin saturation is initialized at equilibrium with the prescribed RBC oxygen partial pressure.

### S8 Additional Results

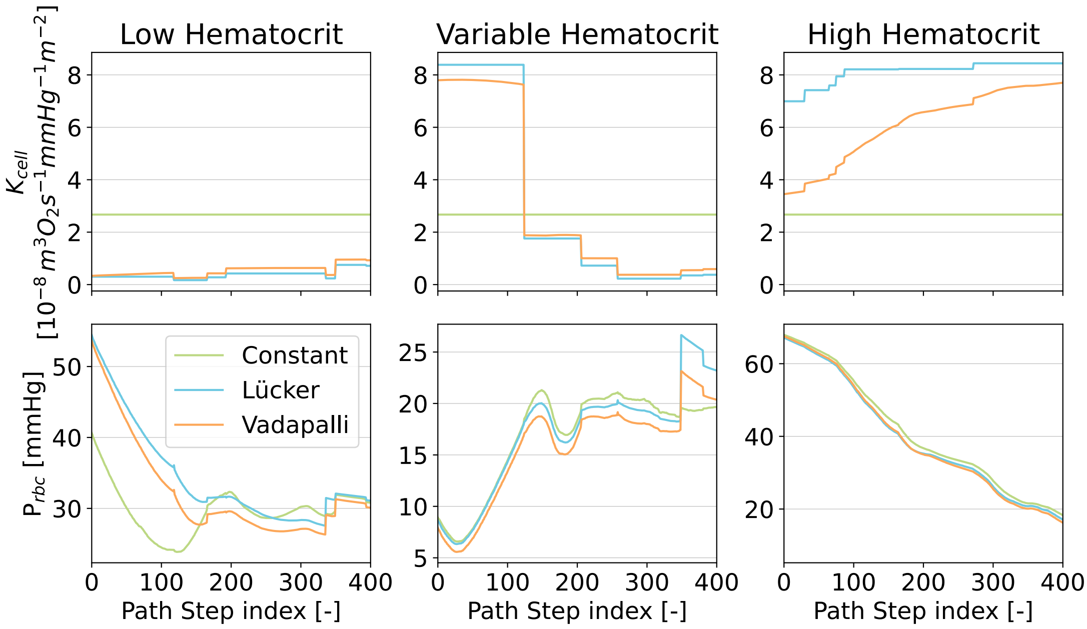


**Fig. S2 Impact of different MTC formulations along representative microvascular flow paths** $K_{\text{cell}}$ (top) and $P_{\text{RBC}}$ (bottom) profiles along three representative flow paths within the microvascular network, depicted in function of the path step index. The path step index denotes the sequential position of each 1D discretization cell along the selected flow path. For each path, results obtained using three different mass transfer coefficient formulations are compared: *Constant* (green), *Lücker* (light blue), and *Vadapalli* (orange).

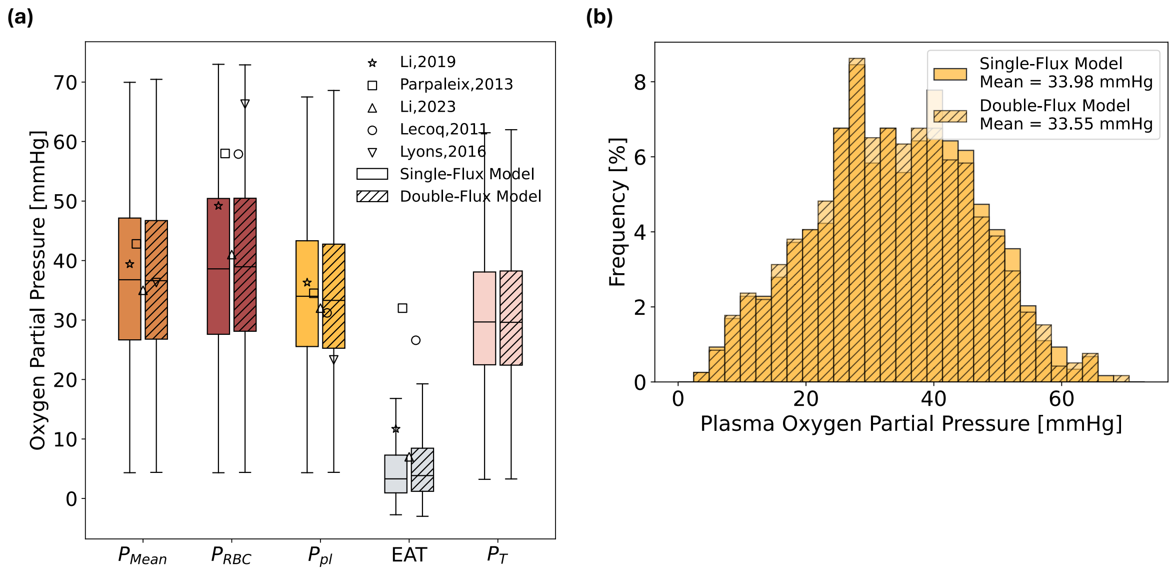


**Fig. S3 Effect of the double-flux formulation on intracapillary and tissue oxygen partial pressure distributions (a)** Boxplots comparing the distributions of mean intracapillary ($P_{Mean}$), red blood cell ($P_{RBC}$), plasma ($P_{pl}$), and tissue ($P_T$) oxygen partial pressure, together with the erythrocyte-associated transient (EAT), obtained with the single- (solid fill) and double-flux model (diagonal hatching). Symbols denote experimental measurements reported in the literature. **(b)** Frequency distributions of $P_{pl}$ across the microvascular network for the single- and double-flux model. Results are shown for a single representative microvascular subnetwork.
